# Visual and semantic representations predict subsequent memory in perceptual and conceptual memory tests

**DOI:** 10.1101/2020.02.11.944801

**Authors:** Simon W. Davis, Benjamin R. Geib, Erik A. Wing, Wei-Chun Wang, Mariam Hovhannisyan, Zachary A. Monge, Roberto Cabeza

## Abstract

It is generally assumed that the encoding of a single event generates multiple memory representations, which contribute differently to subsequent episodic memory. We used fMRI and representational similarity analysis (RSA) to examine how visual and semantic representations predicted subsequent memory for single item encoding (e.g., seeing an orange). Three levels of visual representations corresponding to early, middle, and late visual processing stages were based on a deep neural network. Three levels of semantic representations were based on normative Observed (“is round”), Taxonomic (“is a fruit”), and Encyclopedic features (“is sweet”). We identified brain regions where each representation type predicted later Perceptual Memory, Conceptual Memory, or both (General Memory). Participants encoded objects during fMRI, and then completed both a word-based conceptual and picture-based perceptual memory test. Visual representations predicted subsequent Perceptual Memory in visual cortices, but also facilitated Conceptual and General Memory in more anterior regions. Semantic representations, in turn, predicted Perceptual Memory in visual cortex, Conceptual Memory in the perirhinal and inferior prefrontal cortex, and General Memory in the angular gyrus. These results suggest that the contribution of visual and semantic representations to subsequent memory effects depends on a complex interaction between representation, test type, and storage location.

## Introduction

The term *memory representation* refers to the informational content of the brain alterations that are formed during encoding and recovered during retrieval (Gisquet-Verrier and Riccio 2012; Moscovitch et al. 2016). Given that these brain alterations are hypothesized to consist of persistent synaptic changes among the neurons that processed the original event, the content of memory representations most likely correspond to the nature of analyses performed by the original neurons. In the case of visual memory for objects, these analyses correspond to processing along the ventral (occipitotemporal) pathway from the processing of simple visual features in early visual cortex (e.g., edge processing in V1) to the processing of objects’ identities and categories in anterior temporal and ventral frontal areas (e.g., binding of integrated objects in perirhinal cortex). Thus, object memory representations are likely to consist of a complex mixture of visual and semantic representations stored along the ventral pathway, as well as other regions. Investigating the nature of these representations, and how they contribute to successful memory encoding, is the goal of the current study.

The nature of visual representations has been examined in at least three different domains of cognitive neuroscience: vision, semantic cognition, and episodic memory. Vision researchers have examined the representations of visual properties (Yamins et al. 2014; Rajalingham et al. 2018), semantic cognition researchers, the representations of semantic features and categories (Konkle and Oliva 2012; Clarke et al. 2013; Martin et al. 2018), and episodic memory researchers, the representations that are reactivated during episodic memory tests (Kuhl et al. 2012; Favila et al. 2018). Interactions among these three research domains have not been as intimate as one would hope. The domains of object vision and semantic cognition have been getting closer, both by the fact that semantic cognition researchers often examine the nature of natural object representations at both visual and semantic levels (Devereux et al. 2013; Martin *et al*. 2018), and also that both domains are relying increasingly on the use of advanced neural network models to reveal statistical regularities in object representation (Jozwik et al. 2017; Devereux et al. 2018). However, the episodic memory domain has been somehow disconnected from the other two, partly because it has tended to focus on broad categorical distinctions (e.g., faces vs. scenes) rather than in the component visual or semantic features (Lee et al. 2016). The current study strengthens the links between the three domains by examining how the representations of the visual and semantic features of object pictures predict subsequent performance in episodic perceptual and conceptual memory tasks. When only the visual modality is investigated, the terms “visual” and “semantic” are largely equivalent to the terms “perceptual” and “conceptual,” respectively. To avoid confusion, however, we use the visual/semantic terminology for representations and the perceptual/conceptual terminology for memory tests.

The distinction between perceptual vs. conceptual memory tests has a long history in the explicit and implicit memory literatures, with abundant evidence of dissociations between these two types of tests (for a review, see Roediger and McDermott 1993). Although these dissociations have been typically attributed to different forms of memory *processing* (Roediger et al. 1989) or memory *systems* (Tulving and Schacter 1990), they can be also explained in terms of different memory *representations.* In Bahrick and Boucher’s (1968; 1971) studies, for example, participants encoded object pictures (e.g., a cardinal), and memory for each object was tested twice: first, with a word-based conceptual memory test (*have you encountered a “cardinal”?*), and second, with a picture-based perceptual memory test (*have you seen this particular picture of a cardinal?*). The results showed that participants often remembered the concept of an object but not its picture, and vice versa. The authors hypothesized that during encoding, visual objects generate separate visual and semantic memory representations, and that during retrieval, visual representations differentially contributed to the perceptual memory test, and semantic representations, to the conceptual memory test. This hypothesis aligns with the behavioral principle of transfer appropriate processing (Morris et al. 1977), but expressed in terms of representations rather than forms of processing. In the current study, we investigated the idea of separate visual and semantic memory representations using fMRI and *representational similarity analysis* (RSA).

Although it is common to use a broad distinction between visual and semantic processing/representations in behavioral memory studies (Bahrick and Boucher 1968; 1971; Paivio 1986; Roediger *et al*. 1989), neuroscientists have associated vision and semantics with many different brain regions (e.g., over thirty different visual areas, see Van Essen 2005) and underlying neural signatures. Vision neuroscientists have described the ventral pathway as a posterior-anterior, visuo-semantic gradient, going from occipital regions (e.g. V1-V4), which analyze simple visual features (e.g., orientation, shape, and color), to more anterior ventral/lateral occipitotemporal areas (e.g., lateral occipital complex—LOC), which analyze feature conjunctions, to perirhinal cortex (Barense et al. 2005; Clarke *et al*. 2013) and medial fusiform regions (Martin 2007; Tyler et al. 2013), which analyze integrated objects. Semantic cognition neuroscientists have also supported an anterior-to-posterior analysis progression, often employing RSA or multivoxel pattern analyses (MVPA) to dissociate neural evidence for object categories or properties in this pathway. For example, while taxonomic relationships are commonly reported in the fusiform gyrus and lateral occipital cortex (Mahon et al. 2009; Leshinskaya and Caramazza 2015), more advanced processing of multimodal object properties in perirhinal cortex (Martin *et al*. 2018), the processing of abstract object properties in anterior temporal cortex (Binney et al. 2016), and the control of semantic associations, to the left inferior frontal gyrus (Badre and Wagner 2007) are all consistent with this view. As illustrated by the case of perirhinal cortex, several regions are involved in both visual and semantic processing, which is consistent with the assumption that semantics emerge gradually from vision (Clarke et al. 2015).

If one assumes that memory representations are the residue of visual and semantic processing (Pearson and Kosslyn 2015; Horikawa and Kamitani 2017), then each kind of visual analysis (e.g., processing the color red) or semantic analysis (e.g., identifying a type bird) can be assumed to make a different contribution to subsequent memory (e.g., remembering seeing a cardinal). We can glean information about the representational content of different brain regions by testing whether the evoked representations coded in a region reflect the organizational logic across a set of stimuli along various dimensions, be they perceptual (a region responds similarity to items that are round, or of the same color) or conceptual (a region response more similarity to items whose category structure is similar). These assumptions deserve to be tested, and therefore to investigate the multiplicity of representations and associated brain regions mediating subsequent memory while maintaining parsimony, we focused on only three kinds of visual representations and three semantic representations. For the *visual representations*, we identified three kinds using a *deep convolutional neural network* (DNN) model (Kriegeskorte 2015). DNNs simplify the complexity of visual analyses into a few main kinds, one for each network layer. There is evidence that DNNs can be valid models of ventral visual pathway processing, and can even surpass traditional theoretical models (e.g., HMAX). In the current study, we used three layers of a widely used DNN (VGG16, see Simonyan and Zisserman 2014) to model *Early, Middle,* and *Late* visual analyses (corresponding to the an early input layer, 2^nd^ convolutional layer, and final fully connected layer). For the *semantic representations*, we used three levels that have been previously distinguished in the semantic memory literature (McRae et al. 2005): Observed, Taxonomic, and Encyclopedic features. The *Observed level*, such as “a cardinal is red,” is the closest to vision and comprises verbal descriptions of observable visual features in the presented image. The *Taxonomic level*, such as “a cardinal is bird,” corresponds to a more abstract description based on semantic categories. Although more abstract, this level is still linked to vision because objects belonging to the same category share many visual features (e.g., all birds have two legs which are typically two vertical lines in the visual image). Finally, the *Encyclopedic level,* such as “cardinals live in North and South America,” is the most abstract level because it cannot be typically inferred from visual properties and is usually learned in school or other forms of cultural transmission. Although a model with only three kinds of visual representations and three kinds of semantic representations is an oversimplification, we preferred to start with a simple, parsimonious model, and wait for future studies to add additional or different representation types.

We sought to address the contribution of these various forms of visual and semantic information to episodic memory for object pictures. Our behavioral paradigm consisted of encoding and retrieval phases in subsequent days (**Figure 1**). During encoding, participants viewed pictures of common objects while generating their names, and during retrieval, they performed sequential conceptual and perceptual memory tests. In the *conceptual memory test*, they recognized the names of encoded concepts among the names of new objects of the same categories. In the *perceptual memory test*, they recognized the pictures of encoded objects among new pictures of the same objects (e.g., a similar picture of a cardinal).

**Figure 1.**
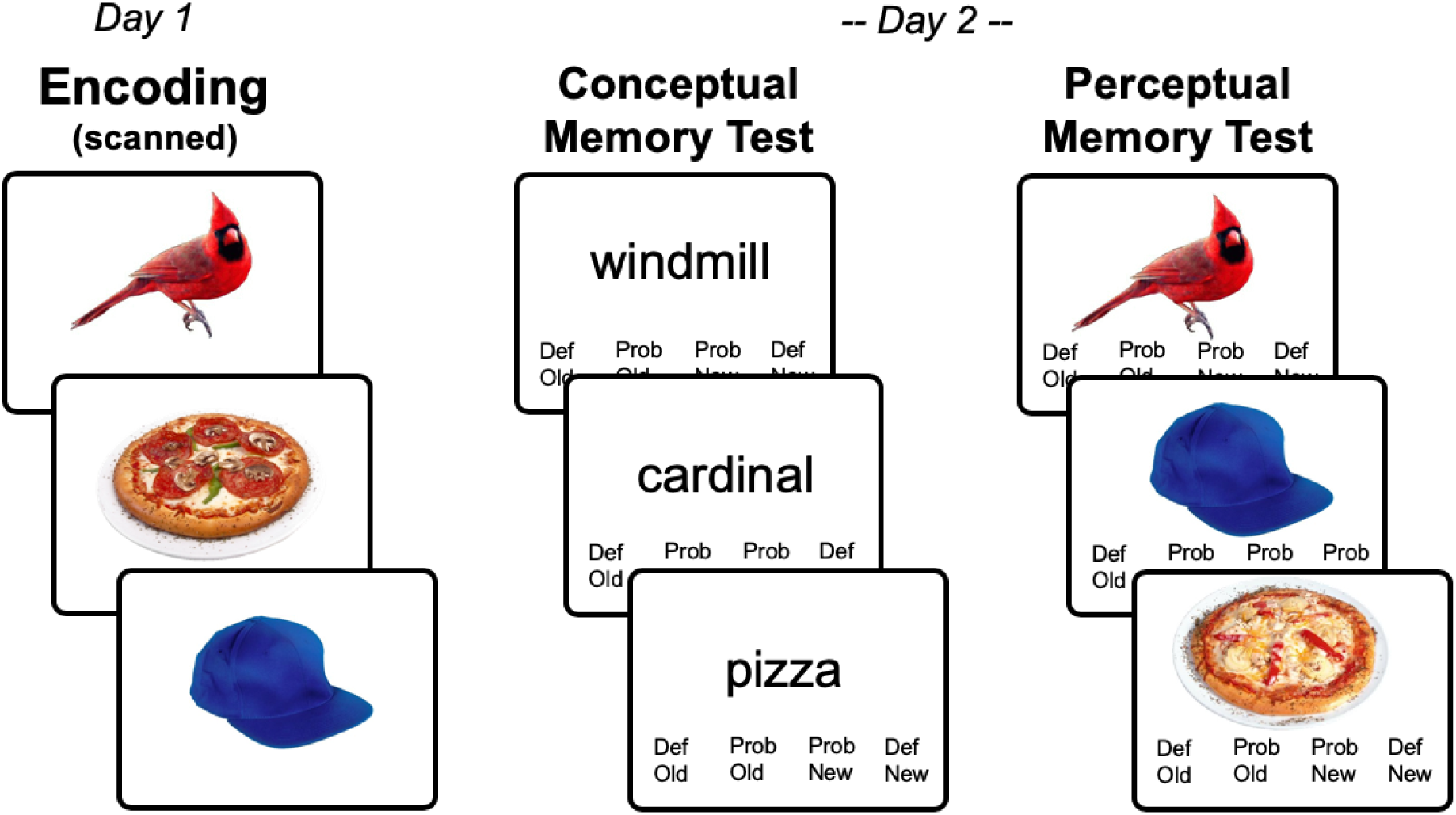
Task Paradigm.

It is important to emphasize that these two tests are not “pure” measures of one kind of memory. We assume that a conceptual memory test is less dependent on the retrieval of visual information than the perceptual. Conversely, we assume that the perceptual memory test is more dependent on visual information than the conceptual memory test because the visual distractors were similar versions of the encoded pictures, and hence, participants had to focus on the visual details to make the old/new decision (in our study we used separate objects, but for an investigation of the effect of object color or orientation on recognition memory, see Brady et al. 2013). However, both conceptual and perceptual tests are also sensitive to the alternative type of information. The conceptual memory test is also sensitive to visual information because participant could recall visual images spontaneously or intentionally to decide if they encountered a type of object. The perceptual memory test is also sensitive to semantic information because different versions of the same object may have small semantic differences that participants may also use to distinguish targets from distractors. Thus, the difference between the informational sensitivity of conceptual and perceptual tests is not absolute but a matter of degree. As a result, we expect some contribution of visual information to the conceptual memory test and of semantic information to the perceptual memory test.

Our method involved four steps (**Figure 2**). The first three steps are standard in RSA studies. (1) The visual and semantic properties of the stimuli were used to create six different *representational dissimilarity matrices* (RDMs). In an RDM, the rows and columns correspond to the stimuli (300 in the current study) and the cell contains values of the dissimilarity (1 – Pearson correlation) between pairs of stimulus representations. The dissimilarity values vary according to the representation type examined. For example, in terms of visual representations, a basketball is similar to a pumpkin but not to a golf club, whereas in terms of semantic representations, the basketball is similar to the golf club but not to the pumpkin. (2) An “activation pattern matrix” was created for each region-of-interest. This matrix has the same structure as the RDM (stimuli in rows and columns) but the cells do not contain a measure of dissimilarity in stimulus properties as in the RDM, but dissimilarity in the fMRI activation patterns by the stimuli. (3) We then computed the correlation between (i) the dissimilarity of *each* object with the rest of the objects in terms of stimulus properties (each row of the RDM) and (ii) the dissimilarity of the same object with the rest of the objects in terms of activation patterns (each row of the activation pattern matrix), and identified brain regions that demonstrated a significant correlation across all items and subjects. We term the strength of this 2^nd^ order correlation the *Itemwise RDM-Activity Fit* (IRAF). The IRAF in a brain region is therefore an index of the sensitivity of that region to that particular kind of visual or semantic representation. Note that such an item-wise approach differs from the typical method of assessing such 2^nd^-order correlations between brain and model RDMs (Kriegeskorte and Kievit 2013), which typically relate the entire item x item matrix at once. This item-wise approach is important for linking visual and semantic representations to subsequent memory for specific objects.

**Figure 2.**
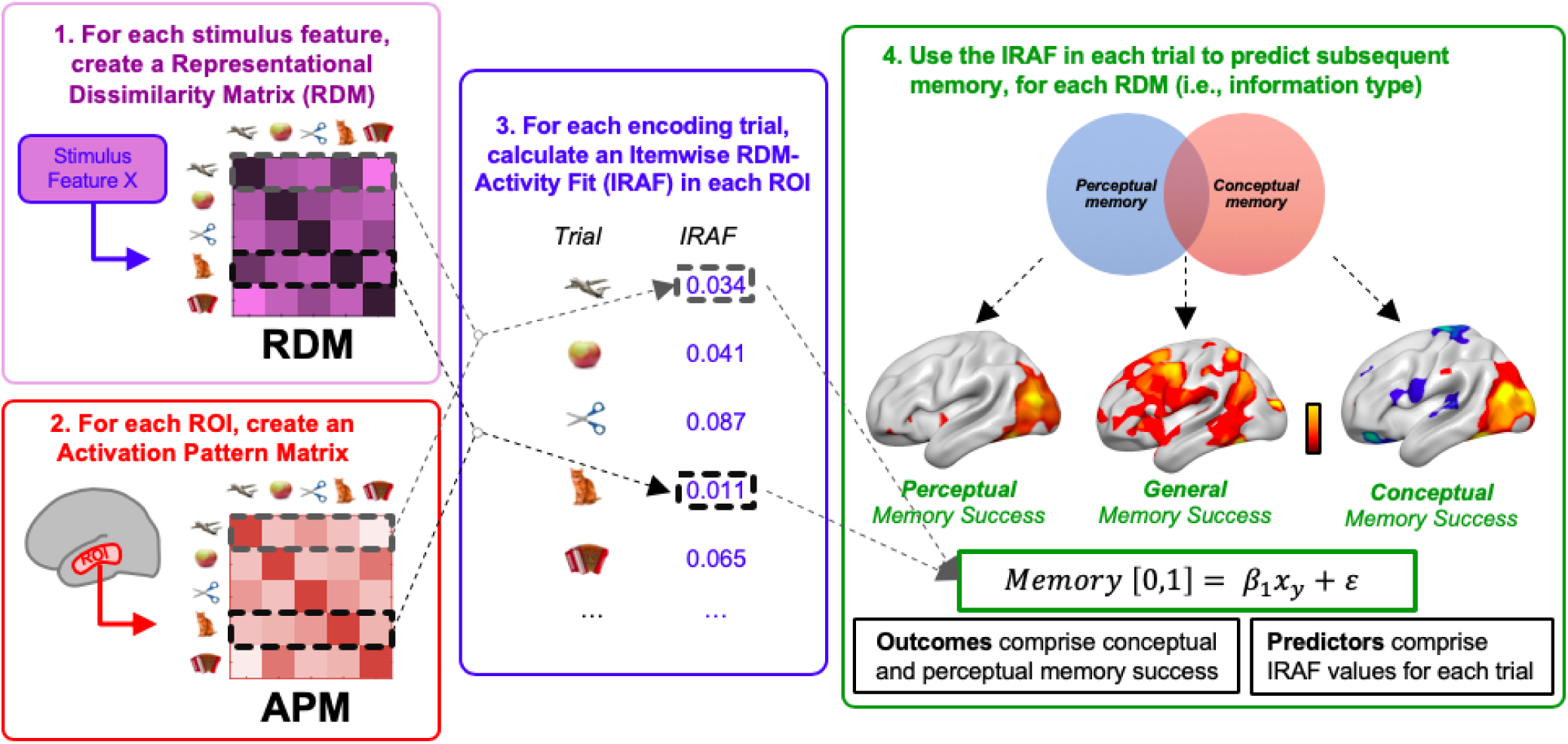
Three steps of the method employed. (1) Representational Dissimilarity Matrices (RDMs) are generated for each visual and semantic representation type investigated, and activation pattern dissimilarity matrices are generated for each region-of-interest. (2) For each brain region, each RDM is correlated with the activation pattern dissimilarity matrix, yielding a Stimulus-Brain Fit (IRAF) measure for the region. (3) The IRAF is used as an independent variable in regressor analyses to identify regions where the IRAF of each RDM predicted subsequent memory in the perceptual memory test but not the conceptual memory test (Perceptual Memory), in the conceptual memory test but not the perceptual memory test (Conceptual memory), and in both memory tests (General Memory).

While the first two steps are standard in RSA studies in the domains of perception (Cichy et al. 2014) and semantics (Clarke and Tyler 2014), and the third is a minor itemwise variation of typical 2^nd^-order brain-model comparisons, our final fourth step is a novel aspect of the current study: (4) we identified regions where the IRAF for each RDM significantly predicted subsequent memory in (1) the perceptual memory test but not the conceptual memory test, (2) in the conceptual memory test but not the perceptual memory test, and (3) in both memory tests (see **Figure 2**). Henceforth, we describe these three outcomes as *Perceptual Memory*, *Conceptual Memory*, and *General Memory.* Furthermore, we expect that regions that are traditionally associated with each of our 6 types of information (and have high IRAF values for their respective models), as well as regions that are not (and have low IRAF values) may both make contributions to subsequent memory in each test. While encoding processes may rely heavily on the visual representations associated with perception (Borst and Kosslyn 2008; Lewis et al. 2011), encoding is not simply a subset of perception, and we therefore expect that such sub-threshold information may nonetheless make a significant contribution to memory strength.

In sum, we extended typical RSA analyses in the domains of vision and semantics by investigating not only what brain regions store different kinds of visual and semantic representations, but also how these various representations predict subsequent episodic memory. Like Bahrick and Boucher (1968; 1971), we hypothesized that visual representations would differentially contribute to the perceptual memory test, and semantic representations, to the conceptual memory test. Given the multiplicity of visual and semantic representations and associated regions, the specific questions we asked is wh re in the brain each kind of visual and semantic representations would differentially predict Perceptual Memory, Conceptual Memory, or General Memory.

## Materials and Methods

### Participants

Twenty-six healthy younger adults were recruited for this study (all native English speakers; 14 females; age mean +/− SD, 20.4 +/− 2.4 years; range 18-26 years) and participated for monetary compensation; informed consent was obtained from all participants under a protocol approved by the Duke Medical School IRB. All procedures and analyses were performed in accordance with IRB guidelines and regulations for experimental testing. Participants had no history of psychiatric or neurological disorders and were not using psychoactive drugs. Of the original participants tested, three participants were excluded due to poor performance/drowsiness during Day 1, one subject suffered a fainting episode within the MR scanner on Day 2, and two participants were subsequently removed from the analysis due to excessive motion, leaving twenty participants in the final analysis.

### Stimuli

Stimuli used in this study were 360 objects drawn from a variety of object categories, including mammals, birds, fruits, vegetables, tools, clothing items, foods, musical instruments, vehicles, furniture items, buildings, and other objects. Additionally, there were also 60 catch-trial items (see behavioral paradigm) evenly distributed from these 12 categories, which were included in the behavioral paradigm but not in the fMRI analyses. During the study, each object was presented alone on white background in the center of the screen with a size of 7.5°.

### Behavioral paradigm

As illustrated by **Figure 1**, the behavioral paradigm consisted of separate encoding and retrieval phases in subsequent days (range = 20-28 hrs, SD = 1.4 hrs). Duren encoding, each trial consisted of an initial fixation cross lasting 500 ms, followed immediately by a single letter probe for 250 ms, immediately followed by an object presented for 500 ms, followed by a blank response screen lasting between 2 and 7 s. Participants we instructed to covertly name each object (e.g., “stork”, “hammer”). In order to ensure they did so, they had to confirm that the letter probe presented before each object matched the object’s name, which was not the case in a small percentage of “catch trials.” If participants could not remember the objects’ name, they pressed a “don’t know” key. Catch trials (10%) and “don’t know” trials (mean=8%) were excluded from the analyses. The object presentation order was counterbalanced across participants, although a constant category proportion was maintained ensuring a relatively even distribution of object categories across the block. This category ordering ensures objects from the 12 different categories do not cluster in time, avoiding potential category clustering as a consequence of temporal proximity. The presentation and timing of stimuli were controlled with Presentation (Psychology Software Tools), and naming accuracy was recorded by the experimenter during acquisition. During retrieval, participants performed sequential conceptual and perceptual memory tests for the encoded objects. In the *Conceptual memory test*, participants recognized the names of encoded concepts among the names of new objects of the same categories. In the *Perceptual memory test*, they recognized the pictures of encoded objects among new pictures of the same objects (e.g., a similar picture of a cardinal). In both tests, participants rated their confidence from definitely new to definitely old.

### MRI Acquisition

The encoding phase and the conceptual memory test were scanned but only the encoding data is reported in this article. Scanning was done in a GE MR 750 3-Tesla scanner (General Electric 3.0 tesla Signa Excite HD short-bore scanner, equipped with an 8-channel head coil). Coplanar functional images were acquired with an 8-channel head coil using an inverse spiral sequence with the following imaging parameters: flip angle = 77°, TR = 2000ms, TE = 31ms, FOV = 24.0 mm^2^, and a slice thickness of 3.8mm, for 37 slices. The diffusion-weighted imaging (DWI) dataset was based on a single-shot EPI sequence (TR = 1700 ms, 50 contiguous slices of 2.0 mm thickness, FOV = 256 × 256 mm2, matrix size 128 × 128, voxel size 2 × 2 × 2 mm, b-value = 1000 s/mm2, 25 diffusion-sensitizing directions, total scan time ∼5 min). The anatomical MRI was acquired using a 3D T1-weighted echo-planar sequence (256 × 256 matrix, TR = 12 ms, TE = 5 ms, FOV = 24 cm, 68 slices, 1.9 mm slice thickness). Scanner noise was reduced with earplugs and head motion was minimized with foam pads. Behavioral responses were recorded with a four-key fiber optic response box (Resonance Technology), and when necessary, vision was corrected using MRI-compatible lenses that matched the distance prescription used by the participant.

Functional preprocessing and data analysis were performed using SPM12 (Wellcome Department of Cognitive Neurology, London, UK) and custom MATLAB scripts. Images were corrected for slice acquisition timing, motion, and linear trend; motion correction was performed by estimating 6 motion parameters and regressing these out of each functional voxel using standard linear regression. Images were then temporally smoothed with a high-pass filter using a 190s cutoff, and normalized to the Montreal Neurological Institute (MNI) stereotaxic space. White matter and CSF signals were also removed from the data, using WM/CSF masks and regressed from the functional data using the same method as the motion parameters. Event-related BOLD responses for correct trials were analyzed using a modified general linear model (Worsley and Friston, 1995) and RSA modelling (described below). Brain images were visualized using the FSLeyes toolbox (fsl.fmrib.ox.ac.uk/fsl/fslwiki/FSLeyes) and SurfIce (www.nitrc.org/projects/surfice/).

### Cortical Parcellation

While voxelwise analyses provide granularity to voxel pattern information they are nonetheless interpreted with respect to specific cortical loci; on the other hand, broad regions of interest (regions-of-interest) encompassing large gyri or regions of cortex often obscure more subtle effects. We chose an intermediate approach and used parcellation scheme with regions of roughly equivalent size and shape. Participants’ T1-weighted images were segmented using SPM12 (www.fil.ion.ucl.ac.uk/spm/software/spm12/), yielding a grey matter (GM) and white matter (WM) mask in the T1 native space for each subject. The entire GM was then parcellated into roughly isometric 388 regions of interest (regions-of-interest), each representing a network node by using a subparcellated version of the Harvard-Oxford Atlas, (Braun et al. 2015), defined originally in MNI space. The T1-weighted image was then nonlinearly normalized to the ICBM152 template in MNI space using fMRIB’s Non-linear Image Registration Tool (FNIRT, FSL, www.fmrib.ox.ac.uk/fsl/). The inverse transformations were applied to the HOA atlas in the MNI space, resulting in native-T1-space GM parcellations for each subject. Then, T1-weighted images were registered to native diffusion space using the participants’ unweighted diffusion image as a target; this transformation matrix was then applied to the GM parcellations above, using FSL’s FLIRT linear registration tool, resulting in a native-diffusion-space parcellation for each subject.

### RSA and subsequent memory analyses

#### 1. Creating representational dissimilarity matrices (RDMs)

An RDM has the stimuli as rows and columns, which each cell indicating the dissimilarity between a pair of stimuli in a particular representation type. For *visual representations*, RDMs were derived from *deep convolutional neural networks* (DNNs; Krizhevsky et al. 2012; LeCun et al. 2015). DNNs consist of layers of convolutional filters and can be trained to classify images into categories with a high level of accuracy. During training, DNNs “learn” convolutional filters in service of classification, where filters from early layers predominately detect lower-level visual features and from late layers, higher-level visual features (Zeiler and Fergus 2014). DNNs provide better models of visual representations in the ventral visual pathway than traditional theoretical models (e.g., HMAX, object-based models; Cadieu et al. 2014; Groen et al. 2018). Therefore, a DNN is an ideal model to investigate multi-level visual feature distinction. Here, we used a pre-trained 16-layer DNN from the Visual Geometry Group, the VGG16 (Simonyan and Zisserman 2014), which was successfully trained to classify 1.8 million objects into 365 categories (Zhou et al. 2017). VGG16 consists of 16 layers including 13 convolutional and three fully-connected layers. Convolutional layers form five groups and each group is followed by a max-pooling layer. The number of feature maps increases from 64, through 128 and 256 until 512 in the last convolutional layers. Within each feature map, the size of the convolutional filter is analogous to the receptive field of a neuron. The trained VGG16 model performance was within normal ranges for object classification (Ren et al. 2017).

We assessed visual information based on the VGG16 model activations from our trained VGG16 model. We used both convolutional (conv) and fully-connected (fc) layers from VGG16. For each convolutional layer of each DNN, we extracted the activations in each feature map for each image, and converted these into one activation vector per feature map. Then, for each pair of images we computed the dissimilarity (squared Euclidean distance) between the activation vectors. This yielded a 300 × 300 RDM for each feature map of each convolutional DNN layer. For each pair of images, we computed the dissimilarity between the activations (squared Euclidean distance; equivalent here to the squared difference between the two activations). This yielded a 300 × 300 RDM for each model unit of each fully-connected DNN layer. We based the *Early visual RDM* in an early input layer, the *Middle visual RDM* in a middle convolutional layer (CV11), and the *Late visual RDM* in the final fully connected layer (FC2).

For the *semantic dimension*, RDMs were based on the semantic features of all observed objects, obtained in a separate normative study (see Supplementary Materials). McCrae feature categories were used to differentiate types of semantic feature information. Feature categories used here include Observed visual features (comprising McCrae feature categories of “visual surface and form”, “visual color”, and “visual-motor”), Taxonomic features, and Encyclopaedic features; feature categories with fewer than 10% of the total feature count (e.g., “smell” or “functional” features) were excluded. The feature vector for the 300 encoding items was 9110 features; objects had an average of 23.4 positive features. The semantic feature RDMs reflect the semantic dissimilarity of individual objects, where dissimilarity values were calculated as the cosine angle between feature vectors of each pair of objects.

The *Observed semantic RDM* was based on the dissimilarity of features that can be observed in the objects (e.g., “is round”), the *Taxonomic semantic RDM* on the dissimilarity on taxonomic or category information (e.g., “is a fruit”), and the *Encyclopedic semantic RDM,* was based on the dissimilarity in encyclopedic details (e.g., “is found in Africa”). It may be worthwhile to note that the Taxonomic semantic RDM, (based on features offered in an independent sample of respondents) has a high overlap with a discrete categorical model based on the explicit category choices listed above (r = 0.92), suggesting that such taxonomic labels faithfully reproduce our a priori category designations.

To provide an intuitive visualization of this novel division of “visual” and “semantic” representations into more granular dimensions, multidimensional scale (MDS) plots for the visual and semantic RDMs are shown in **Figure 3** The MDS plot for the Early visual RDM (**Fig. 3A**) appears to represent largely color saturation (vertical axis) and orientation (horizontal axis); the MDS plot for the Middle visual RDM (**Fig. 3B**) seems to code for shape-orientation combinations (e.g., broad objects at the top-left, thin-oblique objects at the bottom-right); and the MDS plot for the Late visual RDM (**Fig. 3C**) codes for more complex feature combinations that approximate object categories (e.g., animals at the bottom, round colorful fruits to the left, squarish furniture on the top-right). Thus, although we call this RDM “visual” it seems to extends into the semantic dimension. This is not surprising given that the end of the visual processing cascade is assumed to lead into simple semantic distinctions.

**Figure 3.**
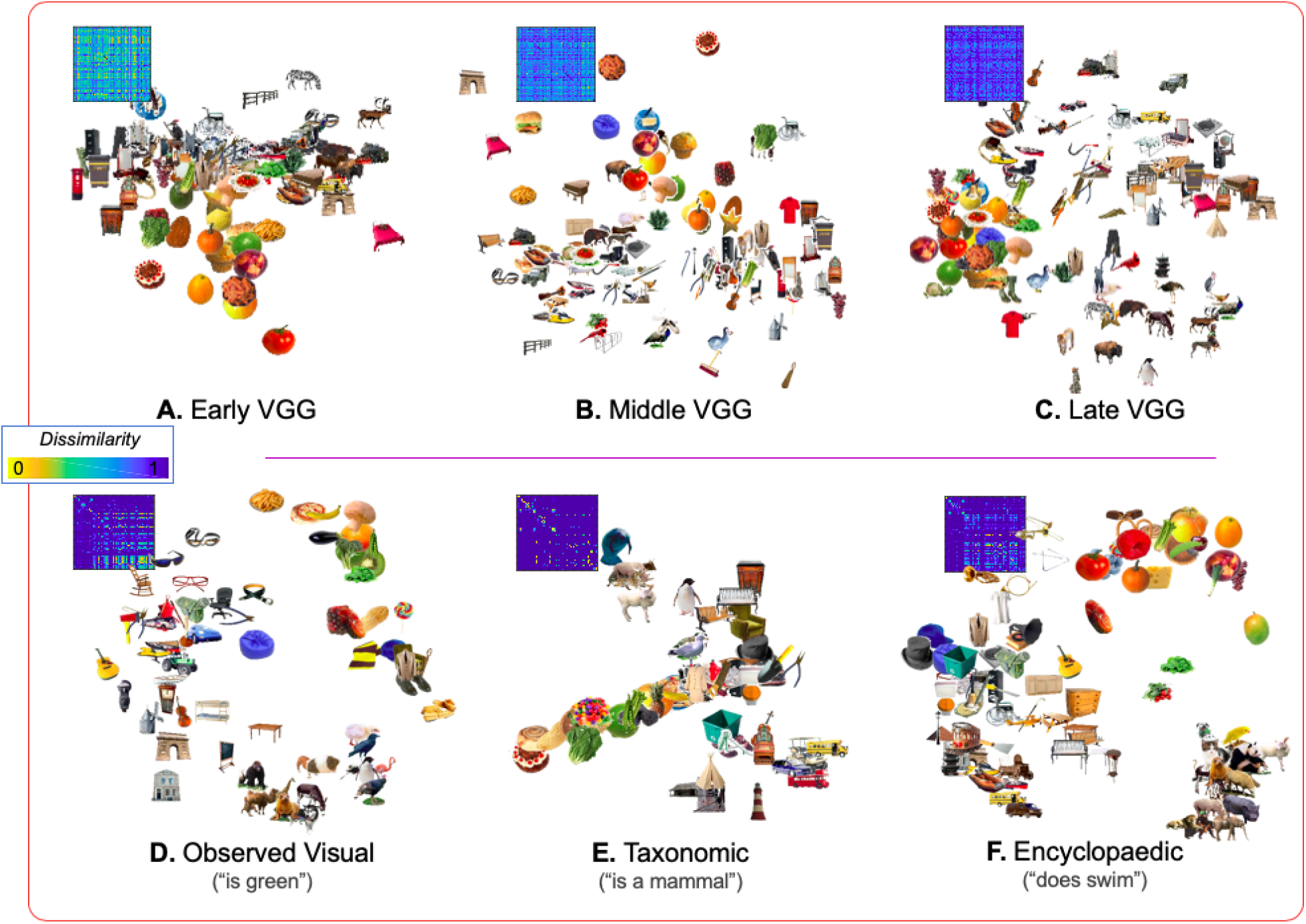
RDMs and corresponding descriptive multi-dimensional scaling (MDS) plots for the three visual and three semantic representations used in the analyses, and which highlight generalized (but by no means complete) dimensions of each type of information. To facilitate representation, we select a subset (n = 80) of the total stimulus set (n = 300) testing in our analyses.

Semantic RDMs displays a similar richness of detail. MDS plot for the Observed semantic RDM (**Fig. 3D**) suggest that this RDM, like the Late visual RDM, codes for the complex combinations of visual features that can distinguish some categories of objects. For example colorful roundish objects (e.g., vegetable and fruits) can be seen on the top right, squarish darker objects (e.g., some furniture and buildings) on the left/bottom-left, and furry animals on the bottom-right. Notably, despite the obvious visual nature of these distinctions, the Observed semantic RDM was created using verbal descriptions of the objects (e.g., “is round”, “is square”, “has fur”) and not the visual properties of the images themselves. The MDS for the Taxonomic visual RDM (**Fig. 3E**), not surprisingly, groups objects into more abstract semantic categories (e.g., edible items to the bottom-left, mammals to the top, and vehicles to the bottom-right). Finally, the MDS for the Encyclopedic visual RDM (**Fig. 3F**) groups objects by features not apparent in the visual appearance (e.g., a guitar and a cabinet appear next to each other possibly because they are made of wood even though their shape is very different). These encyclopedic, non-visual features therefore engender a number of groups that may or may not overlap with the explicit semantic categories used to define our stimulus set. We note also that MDS plots are a largely qualitative, two-dimensional representation of a highlight dimensional feature space (e.g., there are over 2,000 individual Encyclopedic features). An additional, more general qualitative observation is that different visual or semantic features organized object concepts very differently; more quantitatively, this differentiation is captured by the relatively low correlation between all 6 RDMs (all *r* < 0.40). These qualitative and quantitative observations therefore suggest that object representation (and by extension, object *memory*) is not captured by single visual or semantic similarity, but in fact may be more accurately captured by the 6 (and probably more) dimensions used herein.

#### 2. Creating activity pattern matrices

In addition to the model RDMs describing feature dissimilarity, we also created brain RDMs, or activity pattern matrices, which represent the dissimilarity in the voxel activation pattern across all stimuli. Thus, the activation pattern matrices (see **Figure 1**) have a dissimilarity structure as the RDM with stimuli as rows. However, whereas each cell of an RDM contains a measure of dissimilarity in stimulus’ properties, each cell of an activity pattern dissimilarity matrix contains a measure of dissimilarity in activation patterns across stimuli. As noted above, the activation patterns were extracted for 388 isometric regions of interest (ROIs; mean volume = 255 mm^3^), activation values from each region were extracted, vectorized, and correlated across Pearson’s *r*).

#### 3. Identifying brain regions where the RDM-activity fit (IRAF) predict subsequent episodic memory

Each model RDMs was correlated with the activation pattern dissimilarity matrix of each item, in each ROI to obtain a RDM-Activity Fit (IRAF) measure for each item, each region. Spearman’s rank correlations values were Fisher transformed and mapped back to each region-of-interest. Having identified brain regions where each RDM fitted the activity pattern matrix (itemwise RDM-activity fit— IRAF, see **Fig. 2**), we identified regions where the strength IRAF for different RDMs predicted episodic subsequent memory performance. In other words, we used the IRAF as an independent variable in a regression analysis to predict memory in the conceptual and perceptual memory test. Note that such an item-wise approach differs from the typical method of assessing such 2^nd^-order correlations between brain and model RDMs (Kriegeskorte and Kievit 2013; Clarke and Tyler 2014), which typically relate the entire item x item matrix at once, and thus generalize across all items that comprise the matrix, and furthermore do not explicitly assess the model fit or error associated with such a brain-behavior comparison. This more general approach therefore handicaps any attempt to capture both the predictive value of item-specific 2^nd^-order similarity, as well as any attempt to capture the variation of model stimuli as a random effect (Westfall et al. 2016), which we model explicitly below within the context of a mixed-effects logistic model across all subjects and items. Thus, the IRAFs for each visual and semantic RDM were used as predictors in a mixed-effects logistic regression analysis to predict subsequent memory [0,1] for items that were remembered in the conceptual but not the perceptual memory test (Conceptual Memory) vs. items that were remembered in the perceptual but not the conceptual memory test (Perceptual Memory). A separate mixed-effects logistic regression analysis used the same IRAF values to predict items that were remembered in both tests (General Memory). To avoid interactions between visual and semantic representations (or between levels) each region-of-interest was tested independently, and we measure the predictive effect of each model term by examining the t-statistics for the fixed effect for each of the 6 IRAF types (Early, Middle, and Late visual RDMs; Observed, Taxonomic, and Encyclopedic semantic RDMs); subject and stimulus were both entered into both mixed-effects logistic models, but not evaluated further. A generalized R^2^ statistic (Cox-Snell R^2^) was used to evaluate the success of each ROI-wise model, and regions with a model fit below α = 0.05 were excluded from consideration. A FDR correction for multiple comparisons was applied to all significant ROIs, with an effective t-threshold of t = 2.31).

## Results

### Behavioral performance

**Table 1** and **Figure 4** display memory accuracy and response time measures in the conceptual and perceptual memory tests. Hit rates were significantly better for the conceptual than the perceptual memory test (*t*_20_ = 2.60, p = 0.02) but false alarm rates did not differ between them (*t*_20_ = 0.76, p = 0.46). Hits were numerically faster in the conceptual than the perceptual memory test but the difference was not significant in a mixed model (χ^2^= 2.41, p > 0.05). To investigate the dependency between the two tests we used a contingency analysis and the Yule’s Q statistic, which varies from −1.0 to 1.0, with −1 indicating a perfect negative dependency between two measures and 1, perfect positive dependency (Kahana 2000). The Yule’s Q for the conceptual and perceptual memory tasks results was 0.24, indicating a moderate level of independency between the two tests. This finding is consistent with Bahrick and Boucher’s (1968; 1971) findings, and with the assumption that two tests were mediated by partly different memory representations. This result motivates our approach to the brain data, such that we consider the contribution of regional pattern information in predicting memory performance in two separate logistic models, each comprising the diagonals of a memory contingency table: 1) a model predicting items remembered in one test and forgotten in the other, and 2) a model predicting items remembered in both tests from forgotten in both tests.

**Table 1.** Behavioral Performance

| Conceptual Memory | Perceptual Memory |
| --- | --- |

| **Response Accuracy** | **M** | **SD** | **M** | **SD** |
| --- | --- | --- | --- | --- |
| Hit Rate | 0.73 | 0.04 | 0.65 | 0.03 |
| False Alarm Rate | 0.34 | 0.04 | 0.34 | 0.03 |
| d' | 1.09 | 0.14 | 0.84 | 0.12 |
| **Response Time (s)** |  |  |  |  |
| Hits | 1.50 | 0.078 | 1.37 | 0.061 |
| Misses | 1.74 | 0.096 | 1.49 | 0.072 |

**Figure 4.**
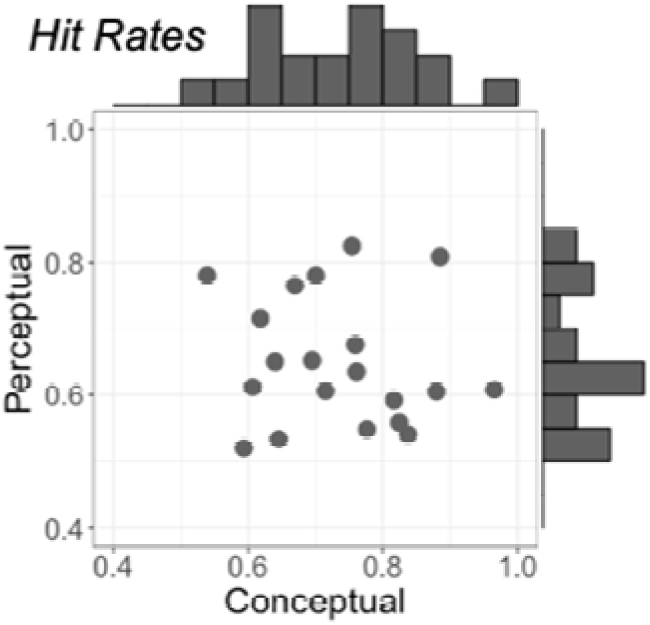
Behavioral Results. Hit rates for both perceptual and conceptual memory tests, plotted by individual.

### Linking RSA to subsequent memory performance

We examined how visual and semantic representations predicted subsequent memory in perceptual and conceptual memory tests. Visual representations were identified using RDMs based on Early, Middle, and Late layers of a deep neural network, and semantic representations using RDMs based on Observed, Taxonomic, and Encyclopedic semantics measures. The Itemwise RDM-Activation Fit (IRAF) was used as a regressor to predict performance for items that were (1) remembered in the perceptual but not the conceptual memory test (Perceptual Memory), (2) remembered in the conceptual but not the perceptual memory test (Conceptual Memory), and (3) remembered in both tests (General Memory). The distribution of IRAF unrelated to memory in these data (see **Fig. S1** for IRAF maps for each of the six RDMs) is generally consistent with both feedforward models of the ventral stream ((early visual RDM showed high IRAF in early visual cortex, later visual RDM representation in more anterior object-responsive cortex, see Konkle and Caramazza 2017), as well as more recent studies focused on the representation of semantic features (e.g., extensive RSA effects in fusiform, anterior temporal, and inferior frontal regions, see Clarke and Tyler 2014). Below, we report regions where IRAFs predicted Perceptual, Conceptual, or General Memory, first for visual RDMs and then for semantic RDMs.

#### Contributions of visual representations to subsequent memory performance

**Table 2** and **Figure 5** show the regions where IRAF in visual RDMs significantly predicted Perceptual, Conceptual, or General Memory, based on a mixed-effects logistic regression analysis in which the information captured in the pattern relationship between fMRI and model dissimilarity (i.e., the IRAF) was used to predict items remembered exclusively either Perceptually or Conceptually, (i.e., the item was remembered in one test but not the other), or a separate logistic model predicting General memory success (i.e. whether a single item was remembered *both* conceptual and perceptual Memory tests).

**Table 2.**
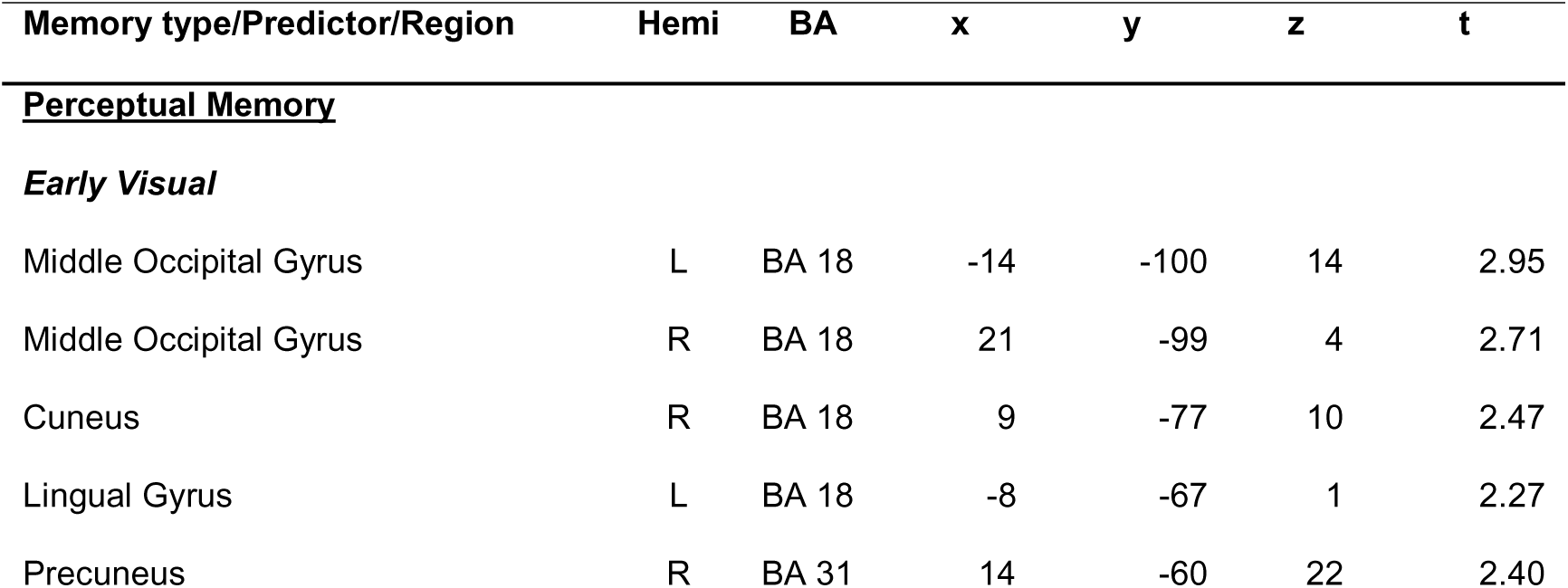
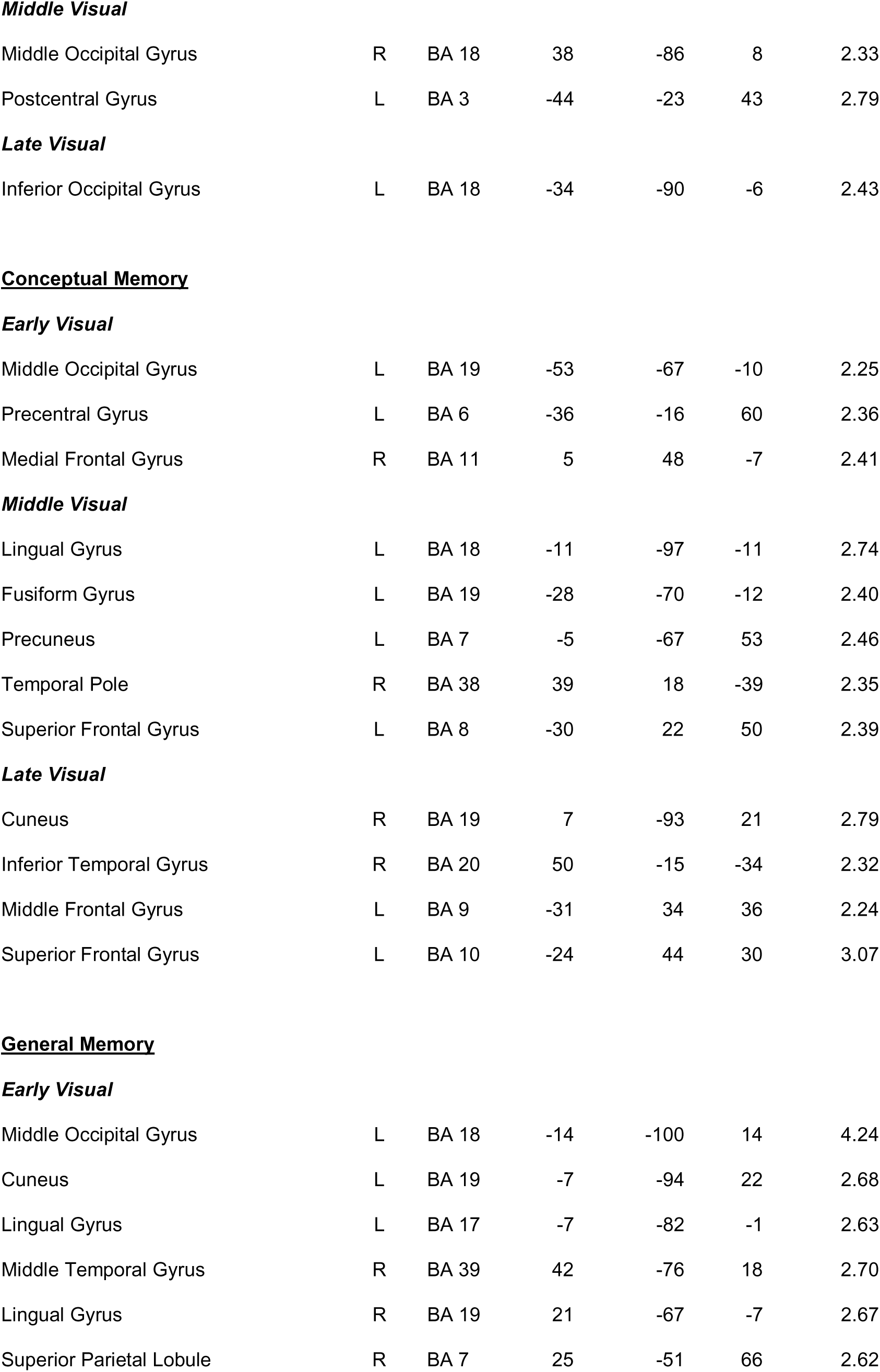
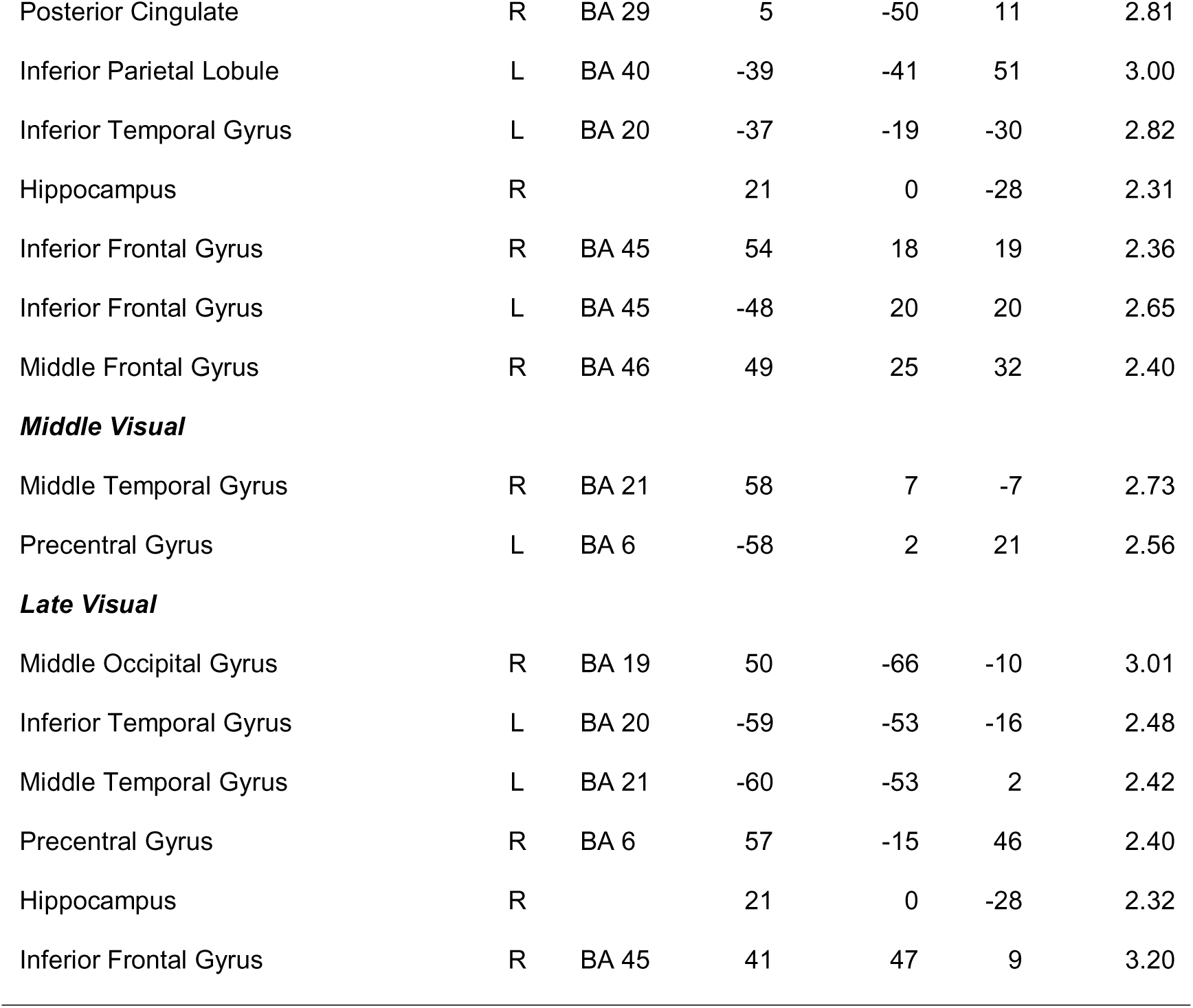
Regions where IRAF values for Early, Middle, and Late visual information predicted Perceptual, Conceptual, or General Memory

| Memory type/Predictor/Region | Hemi | BA | x | y | z | t |
| --- | --- | --- | --- | --- | --- | --- |
| **Perceptual Memory** |  |  |  |  |  |  |
| **Early Visual** |  |  |  |  |  |  |
| Middle Occipital Gyrus | L | BA 18 | -14 | -100 | 14 | 2.95 |
| Middle Occipital Gyrus | R | BA 18 | 21 | -99 | 4 | 2.71 |
| Cuneus | R | BA 18 | 9 | -77 | 10 | 2.47 |
| Lingual Gyrus | L | BA 18 | -8 | -67 | 1 | 2.27 |
| Precuneus | R | BA 31 | 14 | -60 | 22 | 2.40 |
| ***Middle Visual*** |  |  |  |  |  |  |
| Middle Occipital Gyrus | R | BA 18 | 38 | -86 | 8 | 2.33 |
| Postcentral Gyrus | L | BA 3 | -44 | -23 | 43 | 2.79 |
| ***Late Visual*** |  |  |  |  |  |  |
| Inferior Occipital Gyrus | L | BA 18 | -34 | -90 | -6 | 2.43 |
| Middle Occipital Gyrus | L | BA 19 | -53 | -67 | -10 | 2.25 |
| Precentral Gyrus | L | BA 6 | -36 | -16 | 60 | 2.36 |
| Medial Frontal Gyrus | R | BA 11 | 5 | 48 | -7 | 2.41 |
| ***Middle Visual*** |  |  |  |  |  |  |
| Lingual Gyrus | L | BA 18 | -11 | -97 | -11 | 2.74 |
| Fusiform Gyrus | L | BA 19 | -28 | -70 | -12 | 2.40 |
| Precuneus | L | BA 7 | -5 | -67 | 53 | 2.46 |
| Temporal Pole | R | BA 38 | 39 | 18 | -39 | 2.35 |
| Superior Frontal Gyrus | L | BA 8 | -30 | 22 | 50 | 2.39 |
| ***Late Visual*** |  |  |  |  |  |  |
| Cuneus | R | BA 19 | 7 | -93 | 21 | 2.79 |
| Inferior Temporal Gyrus | R | BA 20 | 50 | -15 | -34 | 2.32 |
| Middle Frontal Gyrus | L | BA 9 | -31 | 34 | 36 | 2.24 |
| Superior Frontal Gyrus | L | BA 10 | -24 | 44 | 30 | 3.07 |
| Middle Occipital Gyrus | L | BA 18 | -14 | -100 | 14 | 4.24 |
| Cuneus | L | BA 19 | -7 | -94 | 22 | 2.68 |
| Lingual Gyrus | L | BA 17 | -7 | -82 | -1 | 2.63 |
| Middle Temporal Gyrus | R | BA 39 | 42 | -76 | 18 | 2.70 |
| Lingual Gyrus | R | BA 19 | 21 | -67 | -7 | 2.67 |
| Superior Parietal Lobule | R | BA 7 | 25 | -51 | 66 | 2.62 |
| Posterior Cingulate | R | BA 29 | 5 | -50 | 11 | 2.81 |
| Inferior Parietal Lobule | L | BA 40 | -39 | -41 | 51 | 3.00 |
| Inferior Temporal Gyrus | L | BA 20 | -37 | -19 | -30 | 2.82 |
| Hippocampus | R |  | 21 | 0 | -28 | 2.31 |
| Inferior Frontal Gyrus | R | BA 45 | 54 | 18 | 19 | 2.36 |
| Inferior Frontal Gyrus | L | BA 45 | -48 | 20 | 20 | 2.65 |
| Middle Frontal Gyrus | R | BA 46 | 49 | 25 | 32 | 2.40 |
| ***Middle Visual*** |  |  |  |  |  |  |
| Middle Temporal Gyrus | R | BA 21 | 58 | 7 | -7 | 2.73 |
| Precentral Gyrus | L | BA 6 | -58 | 2 | 21 | 2.56 |
| ***Late Visual*** |  |  |  |  |  |  |
| Middle Occipital Gyrus | R | BA 19 | 50 | -66 | -10 | 3.01 |
| Inferior Temporal Gyrus | L | BA 20 | -59 | -53 | -16 | 2.48 |
| Middle Temporal Gyrus | L | BA 21 | -60 | -53 | 2 | 2.42 |
| Precentral Gyrus | R | BA 6 | 57 | -15 | 46 | 2.40 |
| Hippocampus | R |  | 21 | 0 | -28 | 2.32 |
| Inferior Frontal Gyrus | R | BA 45 | 41 | 47 | 9 | 3.20 |

**Figure 5.**
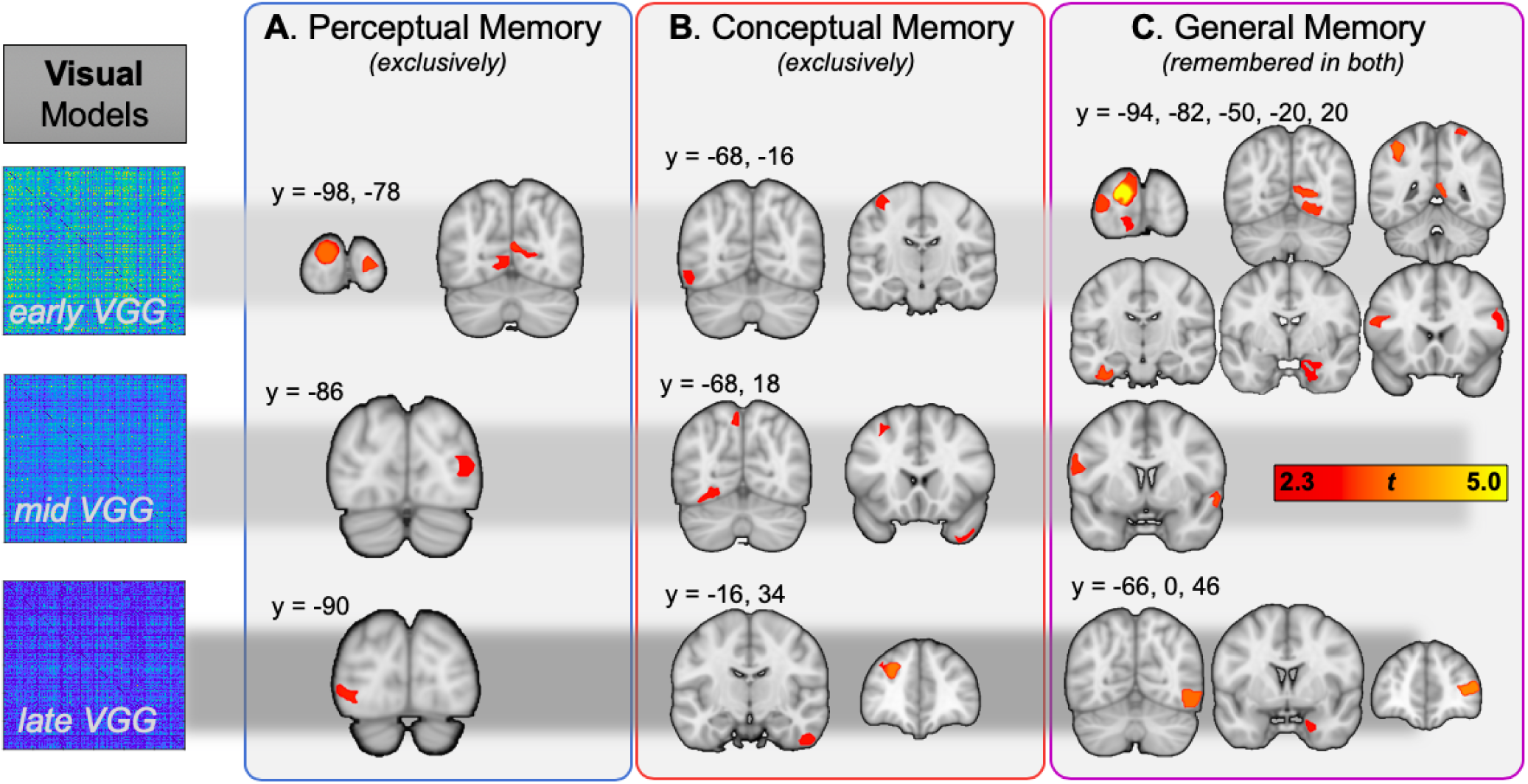
Visual information predicting subsequent Perceptual Memory, Conceptual Memory, and General Memory. The first row represents regions where memory was predicted by Early Visual (Layer 2 from VGG16) information, the second row corresponds to Middle Visual (Layer 12), and the last row to Late Visual (Layer 22) information.

##### Perceptual Memory

The IRAF for the Early Visual RDM predicted Perceptual Memory in multiple early visual regions, in keeping with the expectation that information about basic image properties would selectively benefit subsequent visual recognition. The IRAFs of the Middle and Late Visual RDMs also predicted Perceptual Memory in early visual areas, though these comprise regions further along the ventral stream (generally lateral occipital complex or LOC), and therefore suggest a forward progression in the complexity of visual representations that lead to later perceptual memory. Thus, the visual representations predicting Perceptual Memory (**Figure 5A**) were encoded primarily in visual cortex.

##### Conceptual Memory

In contrast with Perceptual Memory, the visual representations that predicted Conceptual Memory were encoded in more anterior regions (**Figure 5B**). These more anterior regions included the fusiform gyrus, precuneus, and the right temporal pole for the Middle Visual RDM, and lateral temporal cortex and frontal regions for the Late Visual RDM. This result suggests that the effects of memory representations on subsequent memory depend not only on their type but also on where and how they are located in the brain.

##### General Memory

Finally, memory for items that were remembered in both perceptual and conceptual memory tests, or General Memory, was predicted by the IRAFs of visual RDMs in many brain regions (**Figure 5C**). The influence of the early visual RDM was particularly strong, including visual, posterior midline, hippocampal, and frontal regions. In comparison, for visual information based on middle-layer DNN information (layer 12 of the VGG16 model) the right middle temporal gyrus and precentral gyrus made significant contributions to General Memory. Lastly, late visual information (based on the final convolutional layer of our DNN, or layer 22 of the VGG16) was critical for General Memory in lateral occipital cortex (BA19), left inferior and middle temporal gyri, hippocampal, and the right inferior frontal gyrus. The effects in the hippocampus are particularly interesting given the critical role of this structure for episodic memory, and evidence that it is critical it is essential for both perceptual and conceptual memory (Prince et al. 2005; Martin *et al*. 2018; Linde-Domingo et al. 2019).

#### Contributions of semantic representations to subsequent memory performance

Turning to semantic information, we examined how Perceptual, Conceptual, and General Memory were predicted, for each individual trial in each individual, by the three types of semantic information: Observed (e.g., “is round”), Taxonomic (e.g., “is a fruit”) and Encyclopedic (e.g., “is sweet”). The results are shown in **Table 3** and **Figure 6**.

**Figure 6.**
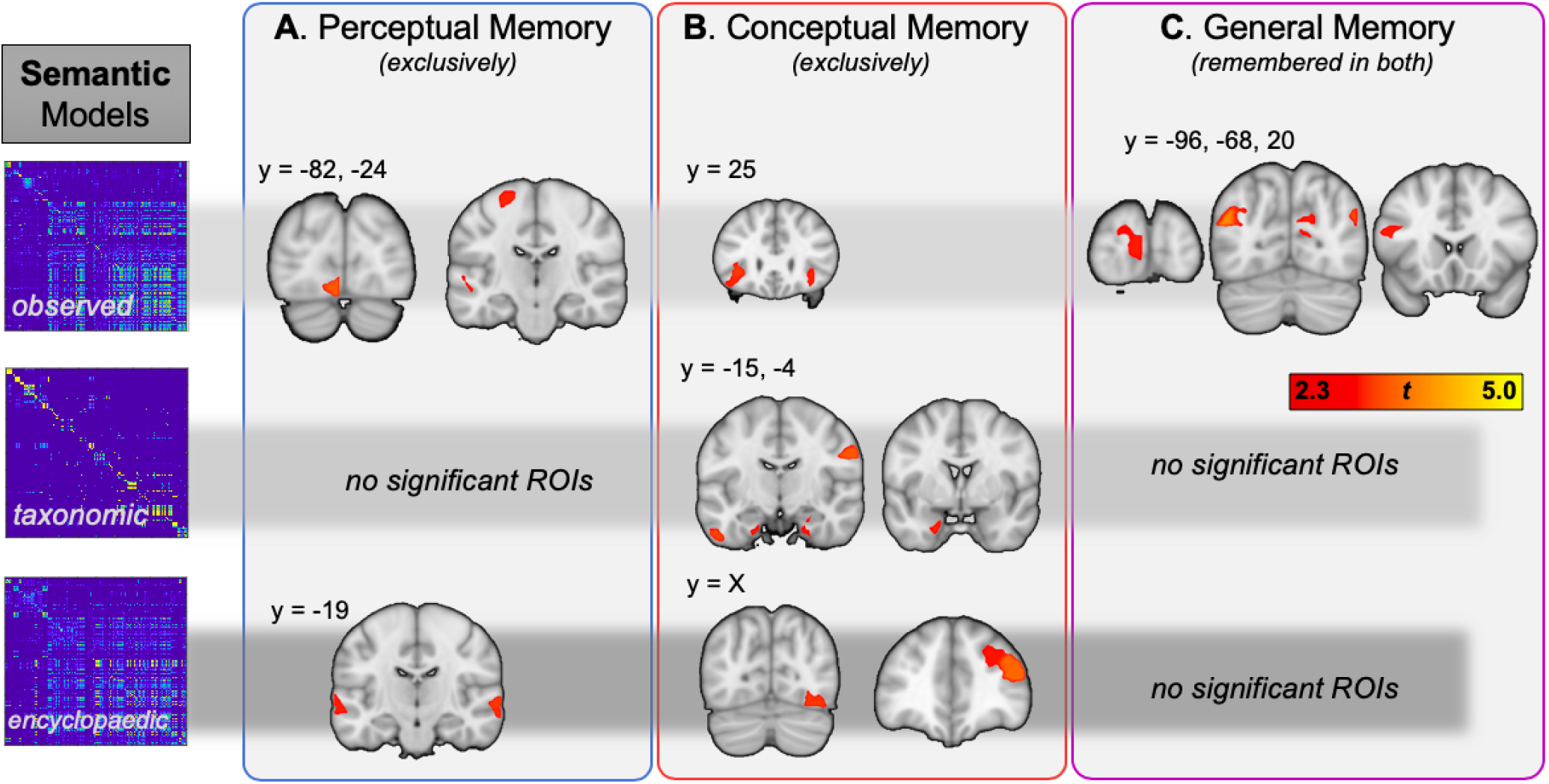
Semantic information predicting subsequent Perceptual Memory, Conceptual Memory, and General Memory. The first row represents regions where memory was predicted by Observed semantic information (e.g., “is yellow”, or “is round”), the second row corresponds to Taxonomic information (e.g., “is an animal”), and the last row to more abstract, Encyclopedic (e.g., “lives in caves”, or “is found in markets”) information.

**Table 3.**
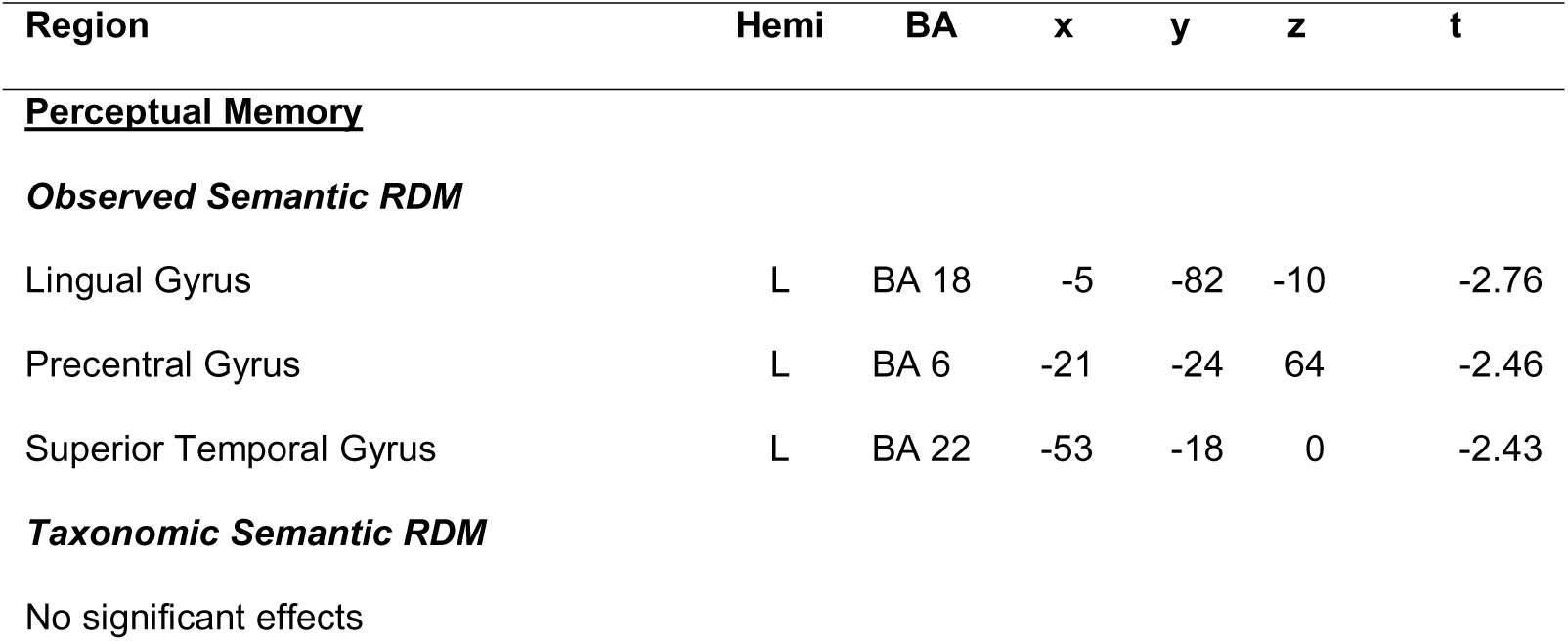
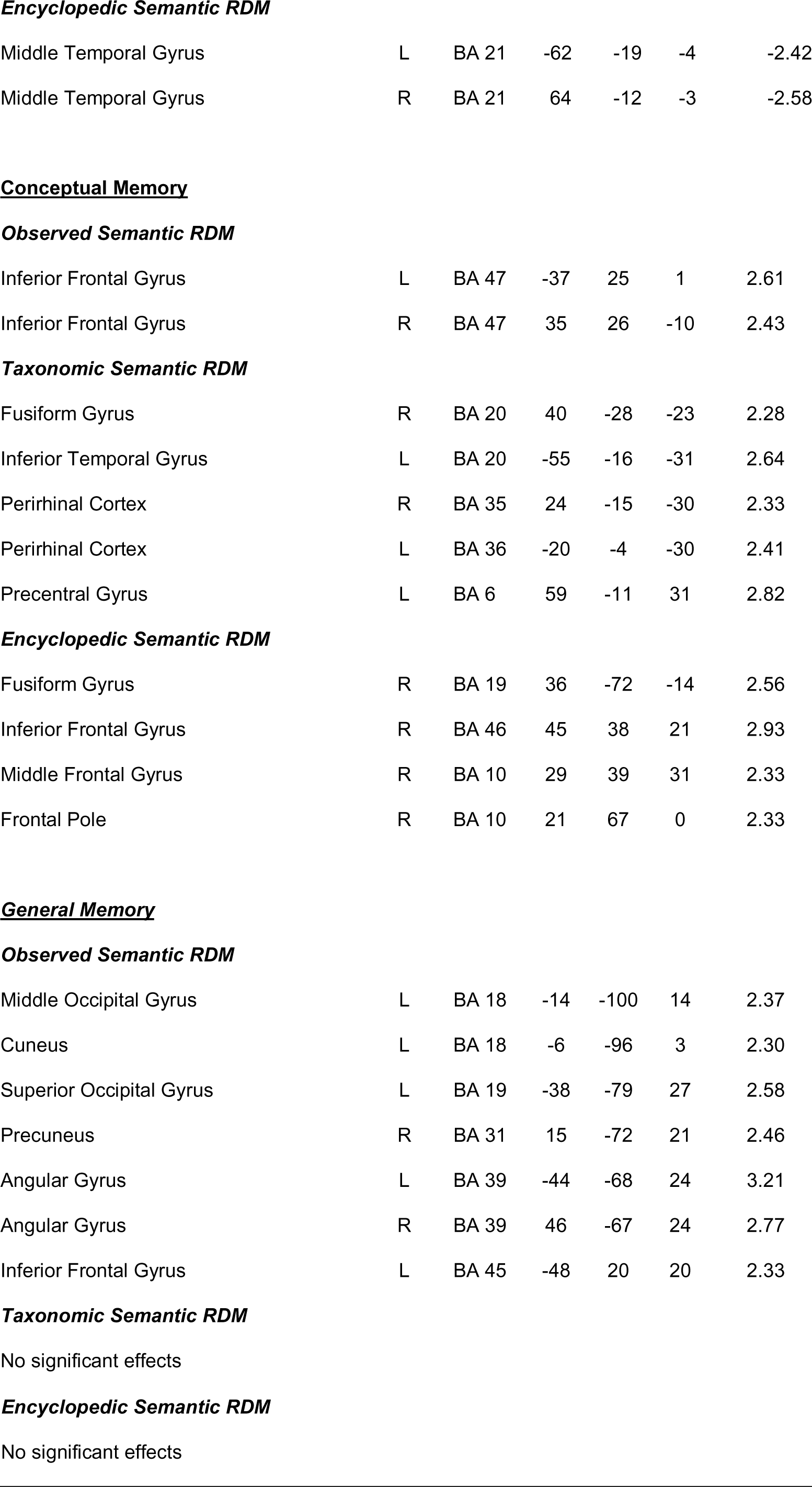
Regions where IRAF values in Observed, Semantic, and Encyclopedic Semantic RDMs predicted Perceptual, Conceptual, or General Memory.

##### Perceptual Memory

Perceptual memory (**Fig. 6A**) was predicted by Observed semantic features stored occipital areas associated with visual processing and left lateral temporal associated with semantic processing. These results are consistent with the fact that Observed semantic features (e.g., “a banana is yellow”) are a combination of visual and semantic properties. Perceptual memory was also predicted by Encyclopedic semantic features (e.g., “bananas grow in tropical climates”) stored in lateral temporal regions. This effect could reflect the role of preexistent knowledge in guiding processing of visual information.

##### Conceptual Memory

Regions where IRF for semantic RDMs predicted Conceptual Memory (**Fig. 6B**) included the left inferior prefrontal cortex for the Observed semantic RDM, the perirhinal cortex for the Taxonomic semantic RDM, and dorsolateral and anterior frontal regions for the Encyclopedic semantic RDM, Taxonomic. The left inferior prefrontal cortex is a region strongly associated with semantic processing, particularly in the anterior are (BA 47) identified (Badre and Wagner 2007). Perirhinal cortex is an area associated with object-level visual processing and basic semantic processing (Tyler *et al*. 2013). Dorsolateral and anterior prefrontal regions are linked to more complex semantic elaboration (Blumenfeld and Ranganath 2007). Thus, conceptual memory was predicted by semantic representations in several regions associated with semantic processing.

##### General Memory

Lastly, memory in both tests (**Fig. 6C**) was predicted only by the IRF of the Observed semantic RDM but not by the IRF of Taxonomic or Encyclopedic RDMs. Given that these two RDMs predicted Conceptual Memory, these results suggest that mid and higher-level semantics contribute specifically to conceptual but not perceptual memory tasks. The regions where Observed semantic representations predicted General Memory included the left inferior prefrontal cortex and the angular gyrus. As mentioned above, the left inferior prefrontal cortex is an area strongly associated with semantic processing (Badre and Wagner 2007). The angular gyrus is also intimately associated with semantic processing (Binder et al. 2009), and there is evidence that representations stored in this region are encoded and then reactivated during retrieval (Kuhl *et al*. 2012).

## Discussion

In the current study we tested the degree to which visual and semantic information could account for the shared and distinct components of conceptual and perceptual memory for real-world objects. We have shown that the encoding of the same object yields multiple memory representations (e.g., different kinds of visual and semantic information) that differentially contribute to successful memory depending on the nature of the retrieval task (e.g., perceptual vs. conceptual). Our results provide compelling evidence that patterns of information commensurate with the retrieval task are beneficial for memory success. Our analysis of selective Perceptual Memory success shows that this form of memory relied on visual information processing in visual cortex, while Conceptual Memory relied on semantic representations in fusiform, perirhinal, and prefrontal cortex—regions typically associated with this form of knowledge representation. However, we also found evidence for a more distributed pattern of mnemonic support not predicted by such a simple view, such that Conceptual Memory benefitted from visual information processing in more anterior regions, (lateral temporal and medial/lateral prefrontal regions), while Perceptual Memory benefitted from semantic feature information processing in occipital and lateral temporal regions. Lastly, General Memory success, which represents strong encoding for items remembered both in the Perceptual and Conceptual memory test, identified differential patterns of regions relying on distinct forms of information, with visual representations predicting General Memory in a large network of regions that included inferior frontal, dorsal parietal, and hippocampal regions, while semantic representations supporting General memory success was expressed in bilateral angular gyrus. The contributions of visual and semantic representations to subsequent memory performance are discussed in separate sections below.

### Contributions of visual representations to subsequent memory performance

Visual representations were identified using RDMs based on a deep neural network (DNN) with three levels of visual representations based on Early, Middle, and Late DNN layers. The use of DNNs to model visual processing along the occipitotemporal (ventral) pathway is becoming increasingly popular in cognitive neuroscience of vision (for review, see Kriegeskorte and Kievit 2013). Several studies have found that the internal hidden neurons (or layers) of DNNs predict a large fraction of the image-driven response variance of brain activity at multiple stages of the ventral visual stream, such that early DNN layers correlate with brain activation patterns predominately in early visual cortex and late layers, with activation patterns in more anterior ventral visual pathway regions (Leeds et al. 2013; Khaligh-Razavi and Kriegeskorte 2014; Güçlü and van Gerven 2015; Kriegeskorte 2015; Wen et al. 2017; Rajalingham *et al*. 2018). Consistent with our results, imagined (versus perceived) visual representations show a greater 2^nd^-order similarity with later (versus early) DNN layers (Horikawa and Kamitani 2017), suggesting a homology between human and machine vision information and their relevance for successful memory formation. However, none of these studies—to our knowledge—have used DNN-related activation patterns to predict subsequent episodic memory.

Notably, many of the regions coding for subsequent memory effects for visual information (**Fig. 5**) lie outside of regions coding for the original perception (**Fig. S1**). Episodic encoding, however, is not simply a subset of perception, and only some information related to a given level of analysis may be encoded, and only a subset of that information may be consolidated. While visual features are likely to be primarily encoded in posterior regions, the representation of those features may be encoded in regions beyond the location of regions traditionally associated with their original encoding, given evidence for color and shape encoding in parietal (Song and Jiang 2006) and prefrontal cortices (Fernandino et al. 2016). The systems consolidation view (Winocur and Moscovitch 2011) purports that such transfer of episodic (from hippocampus to cortex) or working memory traces (from visual to parietal and frontal regions, see Xu 2017). Further evidence from studies scanning both encoding and retrieval blocks, and find that memory representations during retrieval may be found in separate regions from encoding (Xiao et al. 2017; Favila *et al*. 2018) also support this transfer-based view. Furthermore, regions coding for 2^nd^-order similarity between brain and model similarity (i.e., our IRAF regressors) with values below a statistical threshold (and therefore not visible in the IRAF maps in **Fig. S1**) can nonetheless have a highly significant effect on the dependent variable (subsequent memory), as it can happen in any regression analysis. Our results therefore highlight not only the contribution of visual regions to visual information consolidation (a rather limited view), but of any region differentially encoding RDM similarity for a given object.

Like Bahrick and Boucher (1968; 1971), we hypothesized that visual representations would differentially contribute to the perceptual memory test and semantic representations, to the conceptual memory test. However, the results showed that representations predicted subsequent memory depending not only on the kind of representation and the type of test, but also depending on where the representations were located. In the case of visual representations, we found that they predicted Perceptual Memory when located in visual cortex but Conceptual Memory when located in more anterior regions, such as lateral/anterior temporal and medial/lateral prefrontal regions.

We do not have a definitive explanation of why the impact of different representations on subsequent memory depended on their location, but we can provide two hypotheses for further research. A *first hypothesis* is that the nature of representations changes depending on the location where they are stored in ways that cannot be detected with the current RSA analyses. Thus, it is possible that these different aspects of Early visual representations, in the way we defined them, are represented in different brain regions and contribute differently to perceptual and conceptual memory tests. Our *second hypothesis*, which not incompatible with the first, is that representations are the same in the different regions, but that their contributions to subsequent memory vary depending on how a particular region contributes to different brain networks during retrieval. For instance, when Early visual representations are stored in occipital cortex, they might contribute in to a posterior brain network that differentially contributes to subsequent Perceptual Memory, whereas when the same representations are stored in the temporal pole, they might play a role in an anterior brain network that differentially contributes to Conceptual Memory. Such an interpretation would be somewhat inconsistent with a DNN-level interpretation of the visual system, given the 1-to-1 mapping of DNN information to areas of cortex. Furthermore, this hypothesis is not supported by our analysis of the similarity of three DNN-layers independent of memory (first 3 rows of **Fig. S1**), which demonstrate a generally well accepted ventral stream pattern of IRAF values. Nonetheless, our analysis considered each region separately in our memory prediction analysis, it is impossible to discount this network-level hypothesis without a more complex multivariate analysis that would capture representation-level interactions between regions. This interaction between representations and brain networks could be investigated using methods such as representational connectivity analyses (Coutanche and Thompson-Schill 2013).

Visual representations also contributed to performance in *both* memory tests (General Memory). In addition to the same broad areas where visual representations predicted Perceptual and Conceptual Memory, General Memory effects were found in inferior frontal, parietal, and hippocampal regions. Inferior frontal (Spaniol et al. 2009) and parietal, particularly dorsal (Uncapher and Wagner 2009), have been associated with successful encoding, with the former associated with control processes and the latter, with top-down attention processes. The finding that visual representations in the hippocampus contributed to both memory tests is consistent with abundant evidence that this region stores a variety of different representations, including visual, conceptual, spatial and temporal information (Nielson et al. 2015; Mack et al. 2016) and contributes to vivid remembering in many different episodic memory tasks (Kim 2011). Directly relevant to the current study, Prince et al. (2005) investigated univariate activity associated with subsequent memory for both visual (word-font) and semantic (word-word) associations, and though (as in the current study), many regions demonstrated a context-specific manner, such that visual memory success was largely associated with occipital regions, while encoding success for semantic associations was predicted by activity in ventrolateral PFC. However, only one region was associated with memory success for both semantic and perceptual encoding success: the hippocampus, and the confluent findings between this study and our own suggests an important cross-modal similarity between univariate and multivariate measures of encoding success.

### Contributions of semantic representations to subsequent memory performance

As in the case of visual representations, the contributions of semantic representations to subsequent memory depended not only on the test, but also on the locations in which these representations were stored. For example, the semantic representations related Observed semantic features contributed to Perceptual Memory in posterior visual regions, to Conceptual Memory in the left inferior frontal gyrus, and to General Memory in the angular gyrus. These latter brain regions have been strongly associated with semantic processing (Badre and Wagner 2007) and with semantic elaboration during successful episodic encoding (Prince et al. 2007). The finding that Observed semantic representations in the angular gyrus predicted both Perceptual and Conceptual Memory is consistent with the emerging view of this region as the confluence of visual and semantic processing (Binder et al. 2005; Devereux *et al*. 2013). In fact, this region is assumed to bind multimodal visuo-semantic representations (Yazar et al. 2014; Tibon et al. 2019) and to play a role in both visual and semantic tasks (Binder *et al*. 2009; Constantinescu et al. 2016). Although the angular gyrus often shows deactivation during episodic encoding (Daselaar et al. 2009; Huijbers et al. 2012), representational analyses have shown that this region stores representations during episodic encoding that are reactivated during retrieval. For example, Kuhl and colleagues found that within angular gyrus, a multivoxel pattern analysis (MVPA) classifier trained to distinguish between face-word and scene-word trials during encoding, successfully classified these trials during retrieval, even though only the words were presented (Kuhl and Chun 2014).

Taxonomic semantic representations predicted Conceptual Memory in perirhinal cortex. This result is interesting because this region, at the top of the visual processing hierarchy and directly associated with anterior brain regions, is strongly associated with both visual and semantic processing (Clarke and Tyler 2014; Martin *et al*. 2018). However, in our analyses the contribution of this region to subsequent episodic memory was limited to semantic representations, specifically taxonomical, and to the conceptual memory test. The contributions of perirhinal cortex to visual and semantic processing has been linked to the binding of integrated objects representations (Clarke and Tyler 2014; Martin *et al*. 2018). Although integrated object representations clearly play a role in the perceptual memory test, success and failure in this task depends on distinguishing between very similar exemplars, which requires access to individual visual features. Conversely, object-level representations could be more useful during retrieval when only words are provided by the test. This speculative idea could be tested by future research.

Finally, an interesting finding was the fact that General Memory was predicted by Observed semantic representations, but not by Taxonomic and Encyclopedic semantic representations. This result is intuitive when considering that while category-level information is useful in representing an overall hierarchy of knowledge (Connolly et al. 2012) such information provides a rather weak mnemonic cue (“I remember I saw a fruit…”), and such information is typically not sufficient for identifying a specific object. In this case, more distinctive propositional information is necessary for long-term memory (Konkle et al. 2010), and most readily suggested by Observational semantics (“I saw a *yellow* pear”). The utility of concept vs. domain-level naming may shed some light on the utility of general hierarchical knowledge (e.g., taxonomic or category labels) versus more specific conceptual information (e.g., observed details or encyclopedic facts) in predicting later memory. Concepts with relatively more distinctive and more highly correlated distinctive relative to shared features have been shown to facilitate basic-level naming latencies, while concepts with relatively more shared features facilitates domain decisions (Taylor et al. 2012). While drawing a direct link between naming latencies and memory may be somewhat tenuous, the implication is that the organization of some kinds of conceptual information may promote a domain vs. item-level comprehension. Nonetheless, more work must be done at the level of individual items to identify strong item-level predictors of later memory, and this work identifies an item-level technique for relating these kinds of complex conceptual structures with brain pattern information. Furthermore, our findings also challenge the dominant view that regions of the ventral visual pathway exhibit a primary focus for category selectivity (Bracci et al. 2017); given that these models are typically tested with “semantic” information that is comprised of categorical delineations (e.g., faces vs. scenes), it is often unsurprising to find general classification accuracy in fusiform cortex. Nonetheless, when considering either middle Visual information, or Encyclopedic information—both of which demonstrate significant memory prediction scores in our analysis—suggest that such an apparent category selectivity in this region is dependent on both a more basic processing of visual features, as well as more abstract indexing of the abstract Encyclopedic features.

## Conclusion

In sum, the results showed that a broad range of visual and semantic representations predicted distinct forms of episodic memory depending not only on the type of test but also the brain region where the representations were located. Much of our observed results were consistent with the dominant view in the cognitive neuroscience of object vision, that regions in the ventral visual pathway represent a progression of increasingly complex visual representations, and we find multiple pieces of evidence that such visual representations (based on a widely-used DNN model) predicted both Perceptual Memory and General when in primary and extended visual cortices. Furthermore, this visual information made significant contributions to both General and Conceptual Memory when such processing was localized to more anterior regions, (lateral/anterior temporal, medial/lateral prefrontal regions, hippocampus). In turn, semantic representations for Observed visual features predicted Perceptual Memory when located in visual cortex, Conceptual Memory when stored in the left inferior prefrontal cortex, and General Memory when stored in the angular gyrus, among other regions. Taxonomic and Encyclopedic information made contributions limited largely to Conceptual memory, in perirhinal and prefrontal regions, respectively.

This is—to our knowledge—the first evidence of how different kinds of representations contribute to different types of memory test. Broadly, the interesting but unexpected finding that the contributions of visual and semantic representations to perceptual and conceptual memory tests emerge from a such wide diversity of brain regions has important implications for our understanding of representations. Such a set of findings helps to expand our view on the importance of regions outside those typically found in similar RSA analyses of visual properties based on DNN information (Devereux *et al*. 2018; Fleming and Storrs 2019). Thus, many regions appear to be guided by the general principle of transfer-appropriate processing (Morris *et al*. 1977; Park and Rugg 2008), such that congruency between the underlying representational information and the form in which it is being tested leads to successful memory. Nonetheless, we also find evidence for regions outside those traditionally associated with the processing of early or late visual information making significant contributions to lasting representations for objects. We offer two hypotheses for this pattern of findings: first is the hypothesis that the nature of representations varies depending on the region in which they are located, whereas a second—not mutually exclusive—hypothesis is that the nature of representations is constant but what varies is the role they play within brain networks. The first hypothesis could be examined by further examining one kind of representation with multiple RSA analyses and multiple tasks, and the second by connecting representations with brain networks, perhaps via representational connectivity analyses.

## Supporting information

Supplementary Methods and Results

## Acknowledgments

The authors would like to thank Aude Oliva and Wilma Bainbridge for help in the conception of this project, and Alex Clarke for helpful comments on the manuscript.

