## Supplementary Methods and Results for "Visual and semantic representations predict subsequent memory in perceptual and conceptual memory tests"

**Supplementary Materials**

Supplementary Methods

**Semantic Feature Norms**

Semantic similarity values were obtained from a normative study conducted in Amazon Mechanical Turk (AMT), in which 162 AMT workers (18-62 years of age, self-reported native speakers of American English) rated 946 everyday objects (www.cabezalab.org/cabezalabobjects). For each object, the workers selected from a drop-down a connector (*is, has, does, made of*, or “…” for a custom connector) and added an object property (e.g, a canary has wings). Participants were able to take part in repeat sessions and completed between 1 and 5 sessions. Sessions were designed to last about an hour, with 30 concepts presented within each session, and data from 30 participants per concept were used to create a feature x concept production frequency matrix.

Feature responses underwent various stages of processing before their use in the semantic RDM creation, following the procedures used by McRae et al. (2005) and Devereaux et al (2014). These steps included: 1) removal of adverbs, such as *really* and *very*, 2) feature-splitting, for example a feature such as has a round face was rewritten as has a round face and has a face, 3) synonym mapping, which involves identifying synonyms both within and across each concept; for example “*does travel in groups”* and “*does travel in packs”* and “*does travel in a flock”* were collapsed to “*does travel in groups”*, 4) correction of spelling mistakes, 5) morphological mapping, for example “*is used in cooking*” and “*is used by cooks*” were collapsed together as “*is used in cooking*”, 6) removal of plural forms, and 7) removal of features not present in at least two concepts. At all stages in the development and use of these processing rules, results were checked manually and corrected if necessary, to guard against overgeneralized modification of the features. In the last stage of feature processing feature labels from the McCrae database: visual-color (“is blue”), visual-form (“is round”), visual-motion (“does spin”), smell, sound, taste, tactile, function (“is for sleeping”), taxonomic (“is a mammal”), and encyclopedic (“lives in India”). These feature categories were used to classify features by five independent raters with strong inter-rater reliability of features across each raters (ICC > 0.8). After processing, a feature x concept production frequency matrix was created to describe the normalized frequency with a given feature is reported for a given object.

### Supplementary Results

#### Brain regions showing a significant IRAF for each RDM

**Figure S1** and **Table S1** depict regions where the item-wise activation pattern similarity significantly correlated with each of the six RDMs. These results represent the result of a 2^nd^ level t-test across subjects, with the first level representing a t-test across all items (i.e., rows of the RDM corresponding to individual items). As expected, the regions showing a significant IRAF for the Early visual RDM were mainly in early visual cortex, likely including V1 and V2. The regions showing a significant IRAF for the Middle visual RDM were more anterior ventral and lateral regions, possibly including lateral occipital complex (LOC, Eger E et al. 2008). These regions apparently included also V4 (Bartels A and S Zeki 2000), which is not surprising given that the Middle visual RDM was dominated by color information (see **Figure 3**). Lastly, the Late visual RDM correlated with activation patterns in a more widespread set of brain regions, including the bilateral fusiform gyrus, inferior parietal lobule, and the inferior frontal cortex. Taken together, these visual model findings replicate previous feedforward models of a hierarchical visual system (Connolly AC et al. 2012; Clarke A and LK Tyler 2014; Clarke A *et al.* 2015).

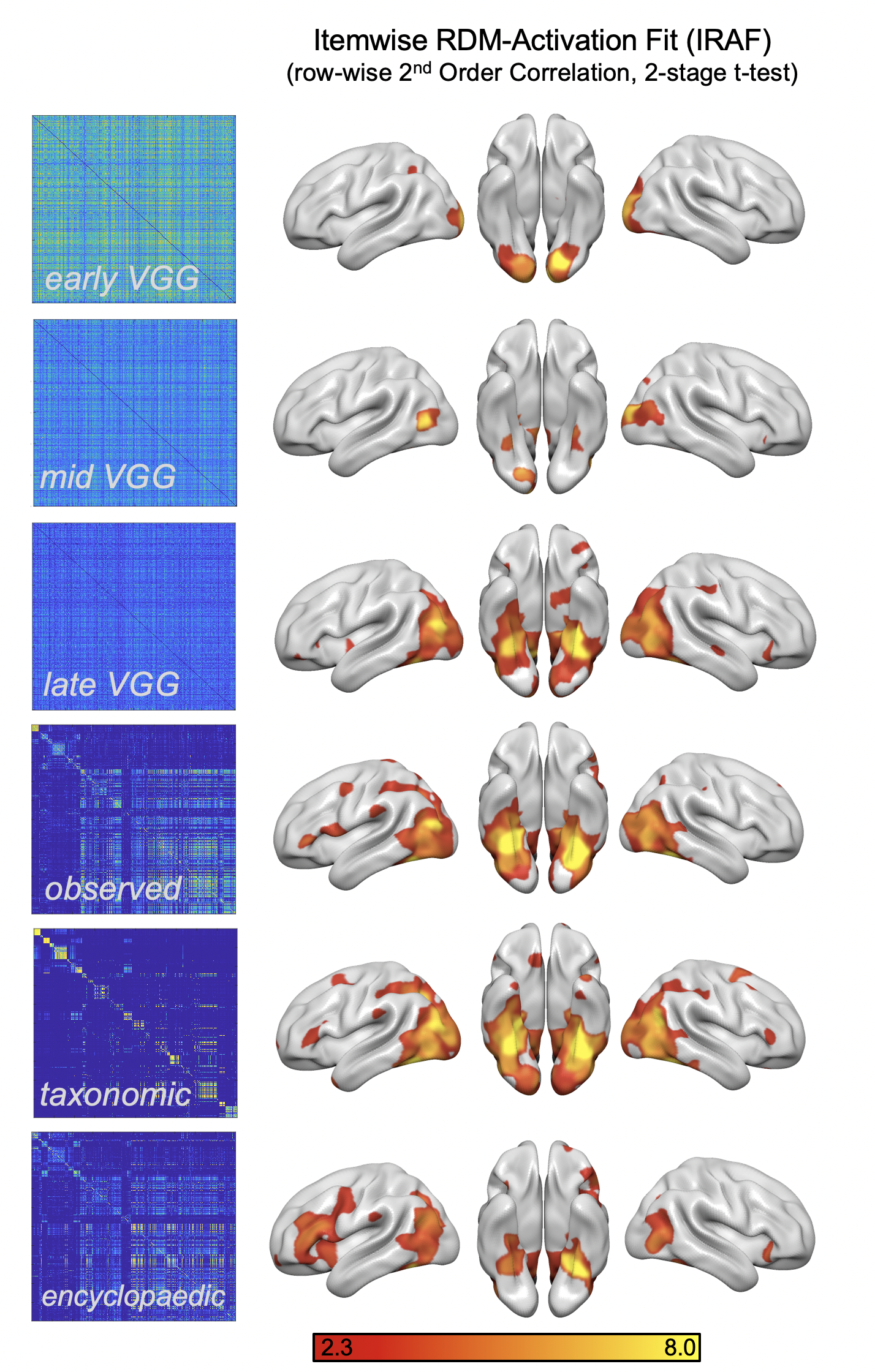

Figure S1. Brain regions where similarity in activation patterns correlated with the similarity in visual or semantic features for each of the six RDMs.

Turning to semantic RDMs, we found that in general, semantic features captured strong item-wise similarity across the entire cortex. Despite an only moderate correlation between the RDM similarity values (all *r* < 0.40), the IRAF patterns for these distinct forms of semantic information showed similar patterns of second-order correlations between brain and model similarity. The Observed and Taxonomic semantic RDMs showed a largely occipitotemporal pattern of IRAF, extending primarily towards ventral anterior temporal regions, as well as a number of smaller prefrontal and parietal regions in the left hemisphere. Finally, the IRAF for the encyclopedic RDM resembles the one for the observed semantic fit but with more limited ventral temporal involvement, and greater brain-model similarity in the left and right prefrontal cortex.

Supplementary Table 1 (S3). Regions associated with each of the six RDMs

| **Model** | **Region** | **Hemi** | **BA** | **x** | **y** | **z** | **t** |
| --- | --- | --- | --- | --- | --- | --- | --- |
| ***VM1*** | *Cuneus* | L | Brodmann area 18/19 | -6 | -96 | 3 | 7.10 |
|  | *Middle Occipital Gyrus* | L | Brodmann area 18 | -19 | -80 | -11 | 6.94 |
|  | *Middle Occipital Gyrus* | R | Brodmann area 18/19 | 21 | -99 | 4 | 6.94 |
|  | *Cuneus* | R | Brodmann area 18 | 13 | -99 | 5 | 6.54 |
|  | *Lingual Gyrus* | L | Brodmann area 18/17 | -5 | -82 | -10 | 5.83 |
|  | *Lingual Gyrus* | R | Brodmann area 17/18 | 6 | -86 | -8 | 5.62 |
|  | *Fusiform Gyrus* | L | Brodmann area 19 | -28 | -70 | -12 | 3.35 |
|  | *Angular Gyrus* | R | Brodmann area 39 | 46 | -67 | 24 | 2.97 |
|  | *Parahippocampal Gyrus* | L | Brodmann area 19 | -23 | -49 | -7 | 2.78 |
|  | *Superior Occipital Gyrus* | L | Brodmann area 19 | -38 | -79 | 27 | 2.76 |
|  | *Middle Temporal Gyrus* | R | Brodmann area 39 | 42 | -76 | 18 | 2.70 |
|  | *Middle Frontal Gyrus* | R | Brodmann area 6 | 24 | 7 | 63 | 2.51 |
|  | *Postcentral Gyrus* | R | Brodmann area 1 | 62 | -24 | 35 | 2.38 |
|  | *Inferior Temporal Gyrus* | R | Brodmann area 37 | 53 | -55 | -7 | 2.37 |
| ***VM2*** | *Cuneus* | R | Brodmann area 18 | 13 | -99 | 5 | 3.96 |
|  | *Middle Temporal Gyrus* | L | Brodmann area 19 | -49 | -70 | 5 | 3.80 |
|  | *Middle Occipital Gyrus* | R | Brodmann area 19 | 26 | -92 | 16 | 3.72 |
|  | *Posterior Hippocampus* | R | Brodmann area 36 | 22 | -34 | -7 | 3.34 |
|  | *Posterior Cingulate* | R | Brodmann area 29 | 5 | -50 | 11 | 3.12 |
|  | *Middle Temporal Gyrus* | R | Brodmann area 39 | 42 | -76 | 18 | 2.92 |
|  | *Lingual Gyrus* | R | Brodmann area 17 | 6 | -86 | -8 | 2.86 |
|  | *Middle Occipital Gyrus* | L | Brodmann area 19 | -44 | -82 | 7 | 2.68 |
|  | *Posterior Cingulate* | L | Brodmann area 30 | -5 | -50 | 14 | 2.66 |
|  | *Inferior Frontal Gyrus* | R | Brodmann area 47 | 35 | 26 | -10 | 2.62 |
|  | *Precuneus* | R | Brodmann area 7 | 21 | -80 | 42 | 2.62 |
| ***VM3*** | *Middle Temporal Gyrus* | L | Brodmann area 19 | -41 | -79 | 15 | 7.77 |
|  | *Precuneus* | R | Brodmann area 31 | 14 | -60 | 22 | 7.21 |
|  | *Parahippocampal Gyrus* | L | Brodmann area 36 | -21 | -37 | -4 | 6.54 |
|  | *Middle Occipital Gyrus* | R | Brodmann area 19 | 46 | -76 | -2 | 6.12 |
|  | *Fusiform Gyrus* | R | Brodmann area 37 | 24 | -47 | -8 | 5.76 |
|  | *Middle Temporal Gyrus* | R | Brodmann area 21 | 42 | -76 | 18 | 5.73 |
|  | *Middle Occipital Gyrus* | L | Brodmann area 19 | -44 | -82 | 7 | 5.48 |
|  | *Superior Occipital Gyrus* | L | Brodmann area 19 | -38 | -79 | 27 | 5.46 |
|  | *Cuneus* | L | Brodmann area 7 | -25 | -77 | 24 | 5.39 |
|  | *Posterior Cingulate* | R | Brodmann area 30 | 16 | -53 | 8 | 5.28 |
|  | *Cuneus* | R | Brodmann area 18 | 13 | -99 | 5 | 5.13 |
|  | *Precuneus* | L | Brodmann area 31 | -13 | -64 | 24 | 4.72 |
|  | *Angular Gyrus* | L | Brodmann area 39 | -44 | -68 | 24 | 4.44 |
|  | *Lingual Gyrus* | L | Brodmann area 18 | -5 | -82 | -10 | 4.09 |
|  | *Superior Parietal Lobule* | R | Brodmann area 7 | 33 | -66 | 39 | 3.74 |
|  | *Superior Parietal Lobule* | L | Brodmann area 7 | -28 | -66 | 40 | 3.70 |
|  | *Posterior Hippocampus* | R | Brodmann area 36 | 22 | -34 | -7 | 3.61 |
|  | *Posterior Cingulate* | L | Brodmann area 30 | -5 | -50 | 14 | 3.56 |
|  | *Perirhinal Cortex* | R | Brodmann area 20 | 33 | -13 | -30 | 3.28 |
|  | *Inferior Frontal Gyrus* | R | Brodmann area 47 | 35 | 26 | -10 | 3.25 |
|  | *Inferior Parietal Lobule* | R | Brodmann area 40 | 54 | -34 | 46 | 3.24 |
|  | *Angular Gyrus* | R | Brodmann area 39 | 46 | -67 | 24 | 3.21 |
|  | *Inferior Temporal Gyrus* | L | Brodmann area 20 | -52 | -55 | -11 | 3.17 |
|  | *Fusiform Gyrus* | L | Brodmann area 19 | -28 | -70 | -12 | 3.16 |
|  | *Hippocampus* | L | Hippocampus | -23 | -5 | -18 | 3.10 |
|  | *Lingual Gyrus* | R | Brodmann area 17 | 6 | -86 | -8 | 2.92 |
|  | *Postcentral Gyrus* | R | Brodmann area 1 | 48 | -21 | 42 | 2.90 |
|  | *Medial Frontal Gyrus* | R | Brodmann area 9 | 7 | 42 | 22 | 2.89 |
|  | *Perirhinal Cortex* | L | Brodmann area 36 | -20 | -4 | -30 | 2.86 |
|  | *Inferior Frontal Gyrus* | L | Brodmann area 47 | -37 | 25 | 1 | 2.73 |
|  | *Inferior Occipital Gyrus* | L | Brodmann area 18 | -34 | -90 | -6 | 2.68 |
|  | *Superior Frontal Gyrus* | L | Brodmann area 11 | -35 | 44 | -13 | 2.67 |
|  | *Insula* | L | Brodmann area 13 | -39 | -1 | -2 | 2.65 |
|  | *Frontal Orbital Cortex* | L | Brodmann area 47 | -40 | 38 | -16 | 2.48 |
|  | *Inferior Temporal Gyrus* | R | Brodmann area 37 | 53 | -55 | -7 | 2.46 |
|  | *Superior Temporal Gyrus* | R | Brodmann area 22 | 54 | -51 | 7 | 2.43 |
| ***SM1*** | *Fusiform Gyrus* | L | Brodmann area 19 | -28 | -70 | -12 | 8.32 |
|  | *Fusiform Gyrus* | R | Brodmann area 37/20 | 24 | -47 | -8 | 7.81 |
|  | *Middle Temporal Gyrus* | L | Brodmann area 19/21 | -41 | -79 | 15 | 7.73 |
|  | *Middle Occipital Gyrus* | R | Brodmann area 19 | 46 | -76 | -2 | 5.94 |
|  | *Precuneus* | R | Brodmann area 7/31 | 12 | -61 | 38 | 5.90 |
|  | *Parahippocampal Gyrus* | L | Brodmann area 36 | -21 | -37 | -4 | 5.76 |
|  | *Parahippocampal Gyrus* | L | Brodmann area 19 | -23 | -49 | -7 | 5.76 |
|  | *Middle Temporal Gyrus* | R | Brodmann area 39/21 | 42 | -76 | 18 | 5.33 |
|  | *Lingual Gyrus* | R | Brodmann area 19 | 21 | -67 | -7 | 5.15 |
|  | *Cuneus* | L | Brodmann area 7 | -25 | -77 | 24 | 5.01 |
|  | *Middle Occipital Gyrus* | L | Brodmann area 19 | -45 | -73 | -7 | 4.91 |
|  | *Angular Gyrus* | R | Brodmann area 39 | 46 | -67 | 24 | 4.79 |
|  | *Inferior Temporal Gyrus* | R | Brodmann area 37 | 53 | -55 | -7 | 4.65 |
|  | *Angular Gyrus* | L | Brodmann area 39 | -44 | -68 | 24 | 4.61 |
|  | *Inferior Frontal Gyrus* | L | Brodmann area 47/45 | -50 | 36 | 0 | 4.60 |
|  | *Inferior Occipital Gyrus* | L | Brodmann area 18 | -38 | -82 | -14 | 4.60 |
|  | *Superior Parietal Lobule* | L | Brodmann area 7 | -28 | -66 | 40 | 4.51 |
|  | *Posterior Cingulate* | L | Brodmann area 30 | -12 | -56 | 9 | 4.45 |
|  | *Hippocampus* | L | Hippocampus | -31 | -25 | -15 | 4.39 |
|  | *Superior Temporal Gyrus* | R | Brodmann area 22 | 54 | -51 | 7 | 3.91 |
|  | *Superior Temporal Gyrus* | L | Brodmann area 22 | -47 | 3 | 8 | 3.77 |
|  | *Lingual Gyrus* | L | Brodmann area 17/19 | -14 | -57 | -4 | 3.76 |
|  | *Precuneus* | L | Brodmann area 7/31 | -13 | -64 | 24 | 3.68 |
|  | *Posterior Cingulate* | R | Brodmann area 30 | 16 | -53 | 8 | 3.58 |
|  | *Middle Frontal Gyrus* | L | Brodmann area 6 | -44 | 0 | 48 | 3.38 |
|  | *Inferior Parietal Lobule* | L | Brodmann area 40 | -60 | -29 | 25 | 3.37 |
|  | *Postcentral Gyrus* | L | Brodmann area 3 | -11 | -40 | 72 | 3.14 |
|  | *Superior Parietal Lobule* | R | Brodmann area 7 | 28 | -64 | 48 | 3.06 |
|  | *Perirhinal Cortex* | R | Brodmann area 35 | 24 | -15 | -30 | 3.05 |
|  | *Lateral Occipital Cortex* | L | Brodmann area 37 | -16 | -65 | 62 | 3.04 |
|  | *Medial Frontal Gyrus* | R | Brodmann area 9 | 7 | 46 | 41 | 3.04 |
|  | *Superior Occipital Gyrus* | L | Brodmann area 19 | -38 | -79 | 27 | 2.92 |
|  | *Cuneus* | R | Brodmann area 18 | 9 | -77 | 10 | 2.78 |
|  | *Postcentral Gyrus* | R | Brodmann area 1 | 48 | -21 | 42 | 2.78 |
|  | *Superior Frontal Gyrus* | L | Brodmann area 8 | -30 | 22 | 50 | 2.78 |
|  | *Posterior Cingulate* | L | Brodmann area 30 | -5 | -50 | 14 | 2.77 |
|  | *Perirhinal Cortex* | R | Brodmann area 20 | 33 | -13 | -30 | 2.67 |
|  | *Precentral Gyrus* | L | Brodmann area 6 | -8 | -24 | 73 | 2.57 |
|  | *Inferior Temporal Gyrus* | L | Brodmann area 37 | -52 | -55 | -11 | 2.55 |
|  | *Anterior Cingulate* | L | Brodmann area 32 | -7 | 42 | 3 | 2.46 |
|  | *Orbital Gyrus* | R | Brodmann area 11 | 9 | 28 | -23 | 2.40 |
| ***SM2*** | *Fusiform Gyrus* | R | Brodmann area 20/37 | 28 | -32 | -15 | 8.60 |
|  | *Middle Temporal Gyrus* | L | Brodmann area 21 | -41 | -79 | 15 | 7.63 |
|  | *Angular Gyrus* | R | Brodmann area 39 | 46 | -67 | 24 | 7.56 |
|  | *Superior Occipital Gyrus* | L | Brodmann area 19 | -38 | -79 | 27 | 6.70 |
|  | *Superior Parietal Lobule* | L | Brodmann area 7 | -28 | -66 | 40 | 6.69 |
|  | *Fusiform Gyrus* | L | Brodmann area 19 | -28 | -70 | -12 | 6.38 |
|  | *Middle Occipital Gyrus* | L | Brodmann area 19 | -45 | -73 | -7 | 6.22 |
|  | *Middle Occipital Gyrus* | R | Brodmann area 19 | 46 | -76 | -2 | 6.14 |
|  | *Hippocampus* | L | Hippocampus | -31 | -25 | -15 | 5.85 |
|  | *Middle Temporal Gyrus* | R | Brodmann area 39 | 42 | -76 | 18 | 5.71 |
|  | *Parahippocampal Gyrus* | L | Brodmann area 36 | -23 | -49 | -7 | 5.39 |
|  | *Perirhinal Cortex* | R | Brodmann area 35 | 33 | -13 | -30 | 5.24 |
|  | *Cuneus* | L | Brodmann area 7/19 | -25 | -77 | 24 | 5.23 |
|  | *Posterior Cingulate* | L | Brodmann area 30 | -12 | -56 | 9 | 5.19 |
|  | *Lingual Gyrus* | R | Brodmann area 17/18 | 6 | -86 | -8 | 5.08 |
|  | *Precuneus* | R | Brodmann area 31 | 14 | -60 | 22 | 5.02 |
|  | *Posterior Cingulate* | R | Brodmann area 30 | 16 | -53 | 8 | 4.92 |
|  | *Inferior Occipital Gyrus* | L | Brodmann area 18 | -38 | -82 | -14 | 4.91 |
|  | *Inferior Parietal Lobule* | L | Brodmann area 40 | -30 | -52 | 47 | 4.83 |
|  | *Anterior Temporal Pole* | L | Brodmann area 37 | -52 | -55 | -11 | 4.81 |
|  | *Lingual Gyrus* | L | Brodmann area 17/19 | -7 | -82 | -1 | 4.79 |
|  | *Cuneus* | R | Brodmann area 18/19 | 13 | -99 | 5 | 4.60 |
|  | *Inferior Parietal Lobule* | R | Brodmann area 40 | 44 | -68 | 43 | 4.54 |
|  | *Inferior Temporal Gyrus* | R | Brodmann area 37 | 53 | -55 | -7 | 4.50 |
|  | *Anterior Temporal Pole* | L | Brodmann area 38 | -38 | 18 | -35 | 4.27 |
|  | *Middle Frontal Gyrus* | R | Brodmann area 6 | 24 | 7 | 63 | 4.22 |
|  | *Posterior Hippocampus* | R | Brodmann area 36 | 22 | -34 | -7 | 4.18 |
|  | *Angular Gyrus* | L | Brodmann area 39 | -44 | -68 | 24 | 3.88 |
|  | *Superior Frontal Gyrus* | L | Brodmann area 8 | -5 | 31 | 46 | 3.69 |
|  | *Superior Frontal Gyrus* | R | Brodmann area 8 | 35 | 18 | 54 | 3.53 |
|  | *Superior Temporal Gyrus* | R | Brodmann area 22 | 54 | -51 | 7 | 3.40 |
|  | *Frontal Pole* | L | Brodmann area 10 | -13 | 69 | -2 | 3.38 |
|  | *Postcentral Gyrus* | L | Brodmann area 3 | -44 | -23 | 43 | 3.37 |
|  | *Superior Parietal Lobule* | R | Brodmann area 7 | 33 | -66 | 39 | 3.29 |
|  | *Inferior Frontal Gyrus* | L | Brodmann area 46/47 | -50 | 36 | 0 | 3.28 |
|  | *Precuneus* | L | Brodmann area 31 | -13 | -64 | 24 | 3.27 |
|  | *Medial Frontal Gyrus* | L | Brodmann area 10 | -6 | 47 | 18 | 3.19 |
|  | *Orbital Gyrus* | R | Brodmann area 11 | 9 | 28 | -23 | 3.13 |
|  | *Cingulate Gyrus* | R | Brodmann area 23 | 4 | -28 | 32 | 3.03 |
|  | *Postcentral Gyrus* | R | Brodmann area 1 | 48 | -21 | 42 | 3.01 |
|  | *Inferior Frontal Gyrus* | R | Brodmann area 11/47 | 52 | 34 | 4 | 2.97 |
|  | *Middle Frontal Gyrus* | L | Brodmann area 6 | -33 | 0 | 56 | 2.95 |
|  | *Inferior Temporal Gyrus* | L | Brodmann area 20 | -59 | -53 | -16 | 2.94 |
|  | *Anterior Temporal Pole* | R | Brodmann area 38 | 27 | 8 | -42 | 2.80 |
|  | *Perirhinal Cortex* | L | Brodmann area 35 | -31 | -5 | -41 | 2.80 |
|  | *Medial Frontal Gyrus* | R | Brodmann area 11 | 3 | 32 | -9 | 2.73 |
|  | *Parahippocampal Gyrus* | R | Brodmann area 36 | 36 | -4 | -42 | 2.63 |
| ***SM3*** | *Parahippocampal Gyrus* | L | Brodmann area 19 | -23 | -49 | -7 | 6.94 |
|  | *Middle Occipital Gyrus* | R | Brodmann area 19 | 46 | -76 | -2 | 6.87 |
|  | *Fusiform Gyrus* | R | Brodmann area 20 | 28 | -32 | -15 | 5.43 |
|  | *Cuneus* | L | Brodmann area 19 | -27 | -92 | 19 | 5.15 |
|  | *Angular Gyrus* | L | Brodmann area 39 | -44 | -68 | 24 | 4.88 |
|  | *Middle Temporal Gyrus* | L | Brodmann area 19 | -41 | -79 | 15 | 4.84 |
|  | *Middle Occipital Gyrus* | L | Brodmann area 19 | -45 | -73 | -7 | 4.75 |
|  | *Middle Temporal Gyrus* | R | Brodmann area 19 | 42 | -76 | 18 | 4.63 |
|  | *Precuneus* | R | Brodmann area 31 | 14 | -60 | 22 | 4.38 |
|  | *Inferior Frontal Gyrus* | L | Brodmann area 46 | -48 | 35 | 9 | 4.21 |
|  | *Angular Gyrus* | R | Brodmann area 39 | 46 | -67 | 24 | 4.20 |
|  | *Superior Occipital Gyrus* | L | Brodmann area 19 | -38 | -79 | 27 | 4.20 |
|  | *Superior Parietal Lobule* | L | Brodmann area 7 | -28 | -66 | 40 | 4.14 |
|  | *Parahippocampal Gyrus* | L | Brodmann area 36 | -21 | -37 | -4 | 3.56 |
|  | *Precuneus* | L | Brodmann area 31 | -13 | -64 | 24 | 3.41 |
|  | *Hippocampus* | L | Hippocampus | -31 | -25 | -15 | 3.33 |
|  | *Posterior Cingulate* | R | Brodmann area 30 | 16 | -53 | 8 | 3.20 |
|  | *Fusiform Gyrus* | L | Brodmann area 19 | -28 | -70 | -12 | 3.09 |
|  | *Inferior Temporal Gyrus* | R | Brodmann area 37 | 53 | -55 | -7 | 3.06 |
|  | *Superior Parietal Lobule* | R | Brodmann area 7 | 33 | -66 | 39 | 3.06 |
|  | *Superior Temporal Gyrus* | R | Brodmann area 22 | 54 | -51 | 7 | 2.94 |
|  | *Middle Frontal Gyrus* | L | Brodmann area 6 | -44 | 0 | 48 | 2.59 |
|  | *Superior Frontal Gyrus* | L | Brodmann area 8 | -5 | 31 | 46 | 2.43 |
|  | *Superior Temporal Gyrus* | L | Brodmann area 22 | -47 | 3 | 8 | 2.38 |
|  | *Caudate* | R | Caudate Body | 14 | 3 | 17 | 2.30 |

*Note: The coordinates reported here indicate the centers of mass identified within each anatomical region. Identification of anatomical regions was confirmed via conversion of MNI coordinates to Talairach coordinates with the mni2tal MATLAB routine (*[*http://www.mrc-cbu.cam.ac.uk/Imaging/mnispace.html*](http://www.mrc-cbu.cam.ac.uk/Imaging/mnispace.html)*).*
